# A plasmodesmata-specific exocyst complex regulates symplastic connectivity by affecting callose turnover

**DOI:** 10.64898/2026.07.02.736176

**Authors:** Edita Janková-Drdová, Samuel Haluška, Tetiana Kalachova, Maria Voloshina, Přemysl Pejchar, Jitka Ortmannová, Eliška Škrabálková, Matěj Drs, Judith García-González, Ivan Kulich, Klára Batystová, Tamara Pečenková, Anzhela Antonova, Anastasiia Zhivaeva, Jiří Šantrůček, Karel Janko, Roman Pleskot, Fatima Cvrčková, Viktor Žárský, Martin Potocký

**Affiliations:** Institute of Experimental Botany of the Czech Academy of Sciences, Rozvojová 263, 165 02 Prague, Czech Republic; Department of Experimental Plant Biology, Faculty of Science, Charles University, Viničná 5, 128 44 Prague, Czech Republic; Department of Biochemistry and Microbiology, Faculty of Food and Biochemical Technology, University of Chemistry and Technology, Technická 5, 166 28 Prague, Czech Republic; Institute of Animal Physiology and Genetics of the Czech Academy of Sciences, Rumburská 89, 277 21 Liběchov, Czech Republic; Department of Biology and Ecology, Faculty of Science, University of Ostrava, Chittussiho 10, 710 00 Ostrava, Czech Republic

**Author notes:** Authors for correspondence: Edita Janková-Drdová, Martin Potocký. Institute of Plant Molecular Biology, Biology Centre of the Czech Academy of Sciences, Branišovská 31, 370 05 České Budějovice, Czech Republic (Ivan Kulich).

## Abstract

Plasmodesmata are intercellular channels that mediate symplastic communication between plant cells. Molecular transport through these channels is critically regulated by dynamic callose deposition and degradation, yet the secretory mechanisms that deliver regulatory components to plasmodesmata remain poorly understood. Here, we identify and characterize a non-canonical plasmodesmata-associated module of the exocyst, an evolutionarily conserved protein complex involved in secretory vesicle tethering and exocytosis. Exocyst subunits EXO70G1, SEC15A, EXO84C, and SEC10A specifically accumulate at plasmodesmata, whereas the canonical exocyst subunits EXO70A1 and SEC8 do not. Genetic and interaction analyses show that EXO70G1 acts as a landmark for recruiting SEC15A and EXO84C to plasmodesmata, revealing a distinct mode of exocyst targeting at these membrane domains. EXO70G1-dependent exocyst targeting to plasmodesmata depends on phosphoinositides and sphingolipids, consistent with the specialized lipid environment of plasmodesmal membranes. Loss of EXO70G1 results in increased callose accumulation and reduced symplastic transport, and strongly enhances developmental defects of a callose-overproducing mutant. In addition, *exo70G1* mutants display enhanced resistance to bacterial pathogen *Pseudomonas syringae,* linking reduced plasmodesmal permeability to anti-bacterial defense. Cross-species analysis further indicates that plasmodesmata association is a derived feature of the EXO70G clade, present in angiosperms but absent from non-angiosperm EXO70 homologs. Together, our findings show that exocyst diversification in plants has generated a specialized trafficking module - plasmodesmata-associated exocyst - that links vesicle delivery to callose homeostasis at plasmodesmata, thereby regulating intercellular communication, development, and immunity.

**Teaser:** A specialized secretion module of the exocyst complex regulates plant cell-to-cell connectivity by controlling callose turnover at plasmodesmata

## INTRODUCTION

The emergence of multicellular organisms created a need for efficient communication between specialized cells, a requirement absent in unicellular systems. In metazoans, this communication is mediated by specialized cell–cell junctions, including gap junctions, adherens junctions, tight junctions, and desmosomes (*1*). In comparison to other multicellular organisms, land plants have evolved many specific characteristics, their rigid cell walls and sessile lifestyle being among the most prominent ones. To meet the special requirement for cellular communication, plasmodesmata emerged as a new cell-cell junction type in Streptophytes, with no homology to any previous structures. Plasmodesmata form highly dynamic membrane-lined channels across the cell wall that enable regulated intercellular exchange of nutrients and signaling molecules not only between neighboring cells, but also among distinct organs via so-called symplastic transport (*2*).

Efficient regulation of transport through plasmodesmata is essential for plant function. A major mechanism controlling symplastic transport involves deposition of callose in the neck region of plasmodesmata, which restricts the transport of molecules, and conversely callose degradation, which restores such transport. Callose accumulation is mediated by the delivery and activity of callose synthases, whereas the delivery and activity of β-1,3-glucanases mediate callose degradation and enhance intercellular connectivity. This dynamic regulation is critical for plant development and responses to environmental stress (*3*). However, despite extensive studies of plasmodesmata structure and function, the secretory mechanisms responsible for delivering these components to plasmodesmata remain poorly understood.

Secretory vesicle trafficking is fundamental for maintaining cellular homeostasis and coordinating growth, development, intercellular communication, and responses to abiotic and biotic stress in plants. Secretory vesicles originating from the endoplasmic reticulum are transported via the Golgi apparatus, and their final tethering to the target plasma membrane is mediated by the hetero-octameric protein complex, the exocyst. Structural and biochemical studies demonstrated that the exocyst is organized into two functional tetrameric modules: module 1, composed of SEC3, SEC5, SEC6, and SEC8 subunits, and module 2, composed of SEC10, SEC15, EXO70, and EXO84 subunits (*4*, *5*). Among these subunits, the SEC3 and EXO70 can directly associate with the plasma membrane through protein-lipid interactions and mediate the targeting of the whole complex to the exocytotic sites, thereby bringing secretory vesicles into proximity with the target membrane prior to SNARE-mediated fusion (*6–8*). Notably, the relative contributions of SEC3 and EXO70 to exocyst targeting differ between eukaryotic kingdoms, with EXO70 becoming dominant in land plants (*4*, *9–11*).

In yeast and animals, most exocyst subunits are encoded by single-copy genes. In contrast, land plants exhibit expansion of exocyst into gene families. Several subunits (e.g., SEC3, SEC5, SEC10, SEC15, and EXO84) have relatively few paralogs (two or three). Crucially, the EXO70 subunit expanded into a large multigene family, containing as many as 23 paralogs in *Arabidopsis thaliana*. The expanded EXO70 paralogs are grouped into three major clades (EXO70.1, EXO70.2, EXO70.3), thought to have undergone functional diversification, putatively gaining specialized roles in trafficking pathways (*10*, *12*, *13*).

The EXO70.1 clade, the closest relative of the opisthokont Exo70, is represented by EXO70A1 and EXO70A2 in *Arabidopsis*. Among these, EXO70A1 is predominantly expressed in sporophytic tissues and together with other exocyst subunits encoded by the paralogs SEC3A, SEC5A, SEC10A, SEC15B, and EXO84B, as well as the single-copy core subunits SEC6 and SEC8, regulates multiple secretory processes, including cell plate formation, cell wall deposition, and recycling of plasma membrane proteins (*14–17*). Owing to both its evolutionary origin and its role in general secretion, this EXO70A1-containing assembly has been termed the canonical exocyst (*4*, *10*). Members of the EXO70.2 clade exhibit pronounced tissue and functional specialization. For example, EXO70B2 and EXO70H1 contribute to defense-related secretion during pathogen attack, whereas EXO70H4 is required for callose deposition during trichome development (*18–21*). The EXO70.3 clade, represented by EXO70G1 and EXO70G2 in *Arabidopsis*, remains the least investigated, and its functional roles are largely unknown.

Here, we provide comprehensive evidence that exocyst protein complex directly localizes to plasmodesmata (PD). Moreover, such a plasmodesmata-associated exocyst (PD-exocyst) has a specific subunit composition, consisting of the module 2 subunits EXO70G1, SEC15A, EXO84C, and SEC10A, together with the module 1 subunit SEC5A. We identify EXO70G1 as a key determinant of plasmodesmata targeting through interactions with specific plasma membrane lipids and recruiting additional exocyst components. At the functional level, the loss of EXO70G1 leads to increased callose accumulation, reduced symplastic transport, and enhanced phenotypic defects in combination with the *cals3-3d* mutation. Consistent with reduced intercellular connectivity, *exo70G1* mutants exhibit increased resistance to *Pseudomonas syringae*. Our data demonstrate that an EXO70G1-targeted exocyst module, with a distinct module 2 subunit composition, regulates callose deposition at plasmodesmata and that this function represents an evolutionarily derived feature of the EXO70G clade.

## RESULTS

### A specific set of exocyst module 2 subunits localizes to the plasmodesmata

During systematic microscopic characterization of GFP-tagged exocyst subunits in *Arabidopsis thaliana*, we found that several subunits from exocyst module 2, namely EXO70G1 (member of the EXO70.3 clade), SEC15A, EXO84C, and SEC10A localized preferentially to immobile puncta at the plasma membrane, resembling the typical distribution of plasmodesmata (PD), across a range of vegetative tissues when expressed under their native or constitutive promoters (Figure 1A, Supplementary Figure S1). In contrast, canonical module 2 subunit EXO70A1 and module 1 core subunit SEC8 exhibited a largely uniform plasma membrane localization (Figure 1A, Supplementary Figure S2), consistent with our previous observations (*4*). Interestingly, another module 1 subunit, SEC5A, showed a punctate localization pattern in above-ground tissues but not in roots (Supplementary Figures S1 and S2).

**Figure 1.**
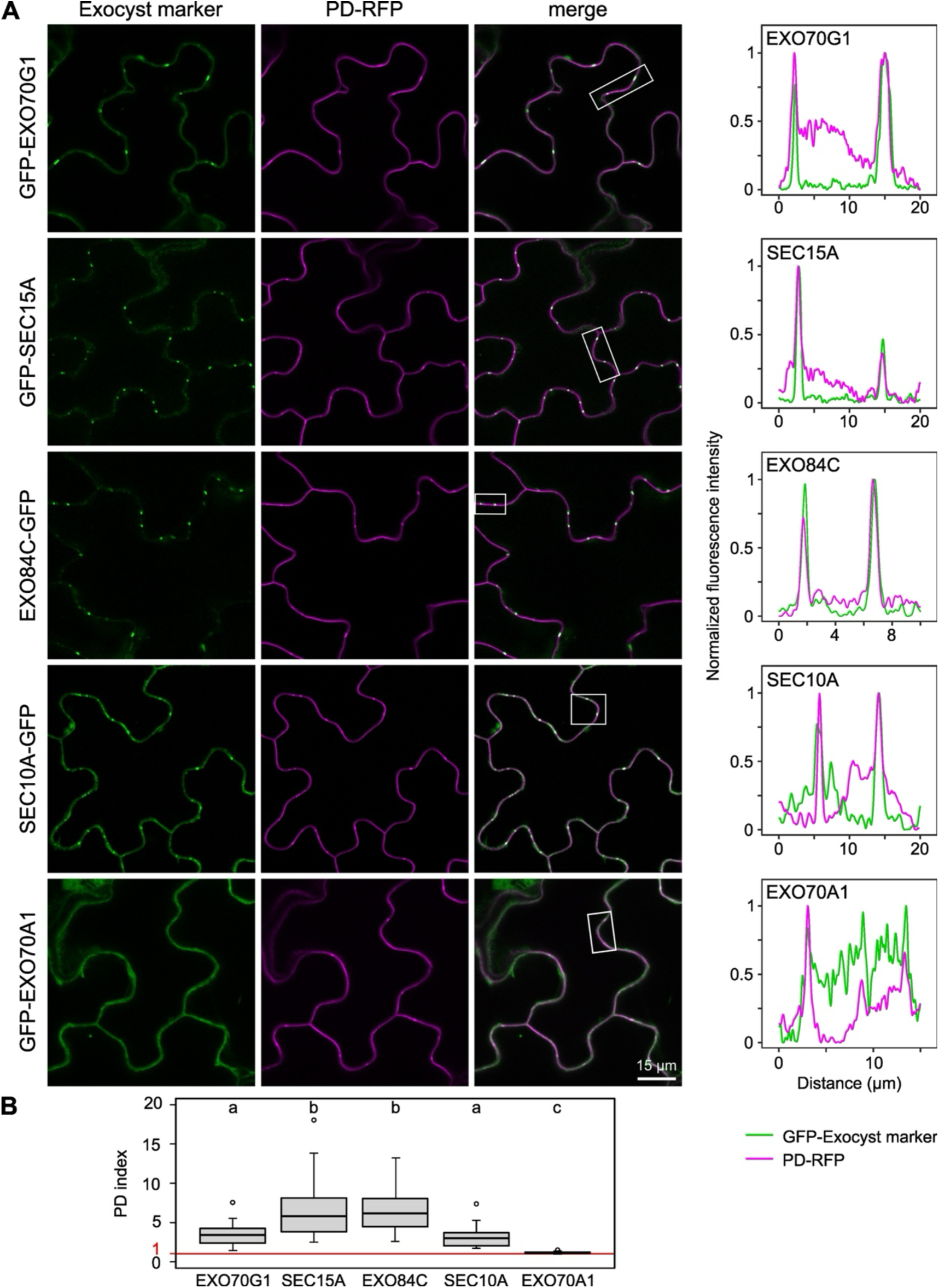
A specific subset of exocyst subunits localizes to plasmodesmata. **(A)** Colocalization of GFP-tagged exocyst subunits EXO70G1, SEC15A, EXO84C, SEC10A, and EXO70A1 with the mRFP-tagged plasmodesmata marker At5g24010 (PD-RFP) in the cotyledon epidermis of 7-day-old transgenic *Arabidopsis thaliana* seedlings. All exocyst subunits were expressed under their native promoters, only EXO70G1 was expressed from 35S promoter. Intensity profiles of GFP and mRFP fluorescence along the plasma membrane are shown for the regions indicated by rectangles. **(B)** Plasmodesmata index (PD index) of the analyzed exocyst subunits, determined from 30 plasmodesmata per sample (five plants). Different letters indicate statistically significant differences (Kruskal-Wallis test, P < 0.05). A PD index value of 1 indicates a uniform distribution of the GFP signal along the plasma membrane.

To confirm that the observed fluorescent puncta correspond to plasmodesmata, we analyzed colocalization of GFP-tagged exocyst subunits with the mRFP-tagged PD marker At5g24010 (hereafter referred to as PD-RFP, (*22*)). Clear colocalization was observed for EXO70G1, SEC15A, EXO84C, and SEC10A (Figure 1A,B), as well as for SEC5A in above-ground tissues (Supplementary Figure S2), but not for EXO70A1 (Figure 1A,B) or SEC8 (Supplementary Figure S2). We independently confirmed the localization of EXO70G1, SEC15A, and EXO84C to plasmodesmata using callose staining by aniline blue fluorochrome (Figure 2).

**Figure 2.**
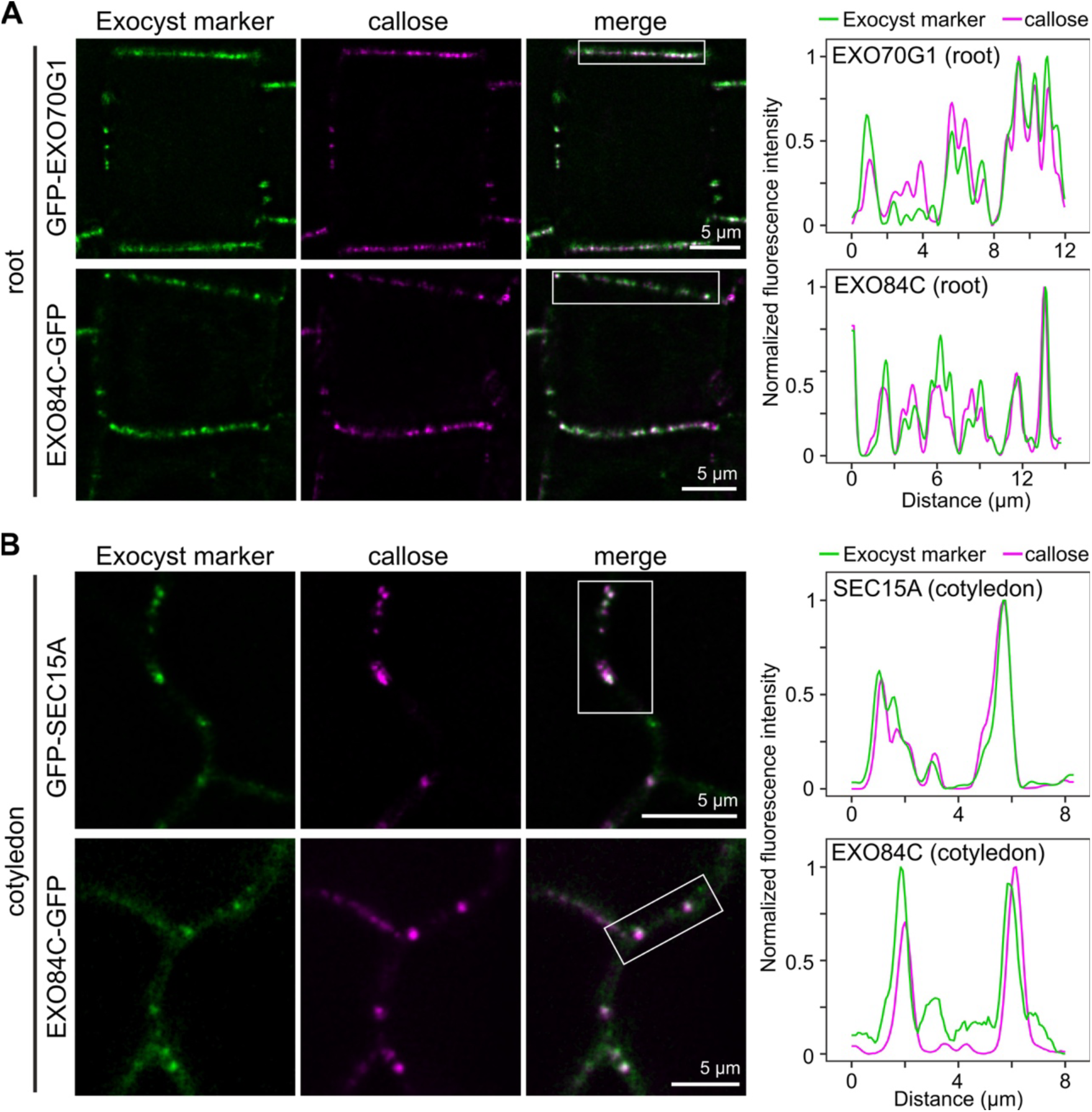
Colocalization of EXO70G1, SEC15A, and EXO84C with callose at plasmodesmata. **(A)** Colocalization of GFP-EXO70G1 and EXO84C-GFP with aniline blue–stained callose in the transition zone of the root epidermis of 7-day-old transgenic *Arabidopsis thaliana* seedlings. **(B)** Colocalization of GFP-SEC15A and EXO84C-GFP with aniline blue–stained callose in the cotyledon epidermis of 7-day-old transgenic *Arabidopsis thaliana* seedlings. Intensity profiles of GFP and aniline blue fluorescence along the plasma membrane are shown for the regions indicated by rectangles in **(A)** and **(B)**. All exocyst subunits were expressed under their native promoters.

To quantify PD association of the tested exocyst subunits, we calculated a PD index, defined as the ratio of GFP fluorescence intensity at PD-RFP-labeled sites to that of the adjacent plasma membrane. This analysis revealed strong enrichment of EXO70G1, SEC15A, EXO84C, and SEC10A and at plasmodesmata (Figure 1B). Moderate enrichment was also observed for SEC5A (median PD index ∼1.8; see Supplementary Figure S2).

In summary, these observations demonstrate that a specific set of exocyst subunits, predominantly from module 2, is enriched at the plasmodesmata. This differential localization pattern points to the existence of a specific plasmodesmata-associated exocyst complex (PD-exocyst), whose composition is distinct from the canonical plasma membrane-localized exocyst complex.

### Module 2 exocyst subunits form plasmodesmata-associated assemblies

Having established that EXO70G1, SEC15A, EXO84C, and SEC10A are enriched at PD, we next examined whether these proteins act as part of a common PD-associated assembly. Consistent with this hypothesis, the colocalization analysis revealed that SEC15A overlaps with EXO84C and SEC10A at plasmodesmata in both leaf and root epidermal cells (Figure 3A, B).

**Figure 3.**
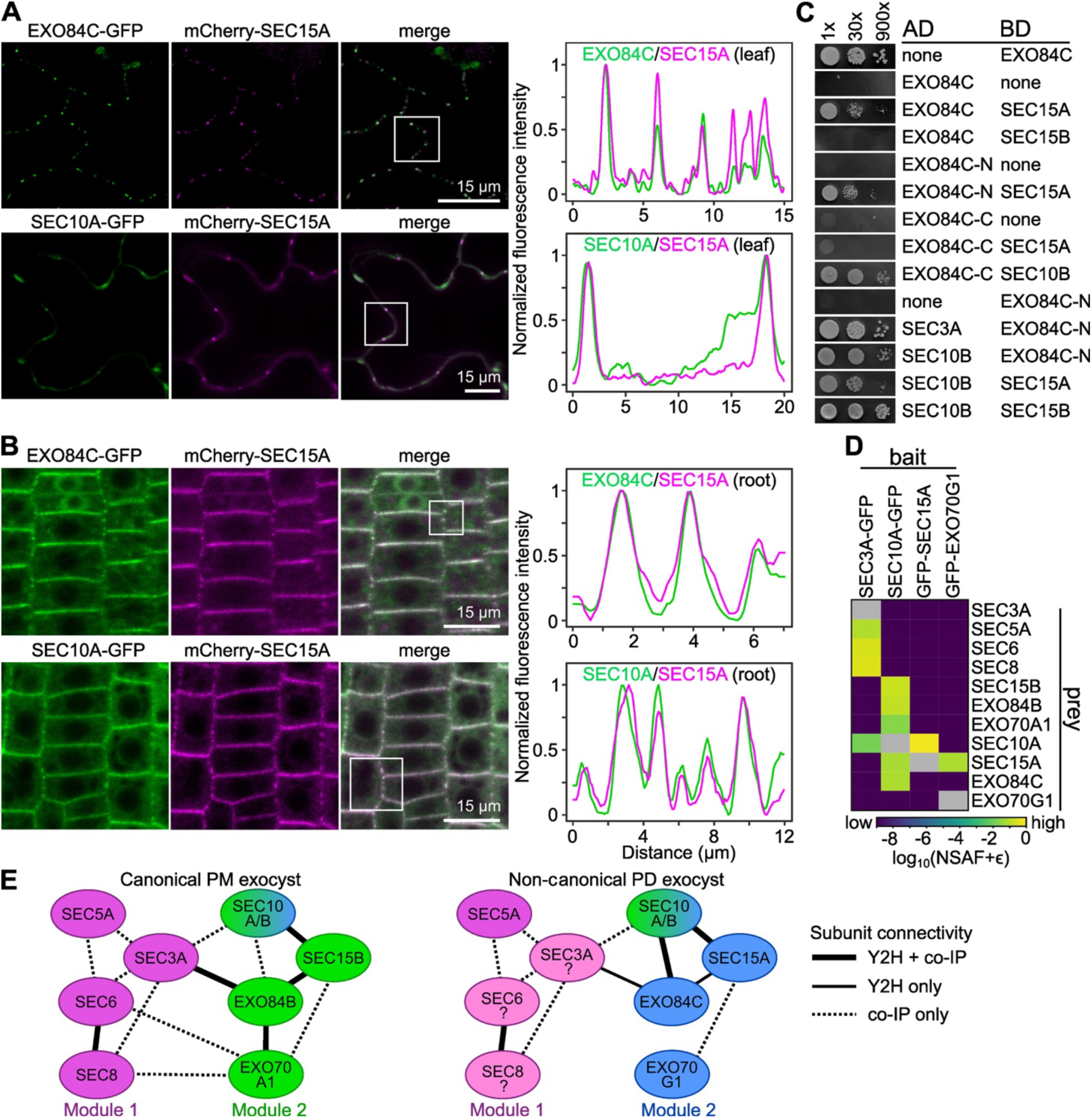
Colocalization and interactions of plasmodesmata-localized exocyst subunits. **(A)** Colocalization of SEC10A-GFP and EXO84C-GFP with mCherry-SEC15A in the leaf epidermis of 10-day-old transgenic *Arabidopsis thaliana* seedlings. **(B)** Colocalization of SEC10A-GFP and EXO84C-GFP with mCherry-SEC15A in the root epidermis of 7-day-old transgenic *Arabidopsis thaliana* seedlings. Intensity profiles of GFP and mCherry fluorescence along the plasma membrane are shown for the regions indicated by rectangles in **(A)** and **(B)**. All tagged exocyst subunits were expressed under their native promoters. **(C)** Interactions among plasmodesmata-localized exocyst subunits and with additional exocyst components, analyzed using the yeast two-hybrid system (AD, activation domain; BD, DNA-binding domain). **(D)** Heatmap showing log₁₀-transformed NSAF (normalized spectral abundance factor) values for exocyst subunits identified by co-immunoprecipitation (Co-IP) with the indicated baits. Color intensity reflects relative protein abundance, with yellow indicating high NSAF values, purple indicating low or background levels, and grey depicting the bait subunit. **(E)** Distinct modular organization and subunit connectivity of canonical PM exocyst (4) and non-canonical PD exocyst complex.

To test preferential pairwise interactions between PD-exocyst subunits, we performed yeast two-hybrid assays. SEC15A, EXO84C, and SEC10B (a near-identical paralog of SEC10A; (*23*)), engaged in mutual interactions (Figure 3C). Notably, full-length EXO84C interacted specifically with SEC15A, the SEC15 paralog enriched at PD, but not with the canonical SEC15 paralog SEC15B, which was reported previously to exhibit a predominantly homogeneous plasma membrane localization (*4*, *24*). Consistent with its conserved role, SEC10B interacted with both PD-localized SEC15A and the PM-localized SEC15B.

Interaction mapping further revealed that both EXO84C C-terminal fragment (EXO84C-C) fused to the Gal4 activation domain (AD) and its N-terminal fragment (EXO84C-N) fused to the Gal4 DNA-binding domain (BD) interact with SEC10B. In contrast, only the N-terminal fragment interacts with SEC15A. Full-length EXO84C fused to the BD could not be analyzed due to autoactivation of reporter gene expression. In addition, the N-terminal EXO84C fragment interacted with the module 1 subunit SEC3A (Figure 3C).

To further investigate the interactome of PD-localized exocyst subunits *in planta*, we performed a series of co-immunoprecipitations of proteins associated with GFP-tagged SEC3A, SEC10A, SEC15A, and EXO70G1, followed by mass spectrometry–based identification. Focusing on exocyst subunits, we found that EXO70G1 associates with complexes containing SEC15A, while SEC15A co-immunoprecipitated with SEC10A. In turn, SEC10A was associated not only with PD-exocyst subunits EXO84C and SEC15A, but also with canonical PM exocyst subunits SEC15B, EXO84B, and EXO70A1. Finally, GFP-tagged SEC3A co-immunoprecipitated core exocyst components SEC5, SEC6, SEC8, as well as SEC10A (Figure 3D; (*4*)).

Together, our colocalization, pairwise interaction, and co-immunoprecipitation data establish a network of direct and indirect interactions between SEC15A, EXO84C, SEC10A/B and EXO70G1, strongly supporting the existence of PD-associated exocyst subunit assembly centered on distinct module 2 subunits (Figure 3E). Whereas SEC10 can be incorporated into both PD-specific and canonical PM exocyst complexes, the preferential interaction of EXO84C with SEC15A over SEC15B may represent a defining feature of the PD-exocyst.

### Recruitment of exocyst subunits to the plasmodesmata depends on EXO70G1

The *Arabidopsis* EXO70A1 subunit has been shown to act as a landmark for the recruitment of canonical exocyst subunits, including SEC6, SEC8, SEC10A/B, SEC15B, and EXO84B, to the lateral membrane of root tip cells (*4*). We therefore asked whether EXO70G1 or other PD-exocyst subunits similarly recruit additional exocyst components to plasmodesmata.

To address this question, we expressed GFP-tagged EXO70G1, SEC15A, and EXO84C in *exo70G1*, *sec15A*, and *exo84C* loss-of-function backgrounds. For comparison, we included *exo70A1* and *exo84B* mutants, which are defective in canonical PM exocyst subunits (*4*, *14*). Notably, none of the PD-localized subunit mutants exhibit any obvious sporophytic defect ((*24*, *25*); Supplementary Figures S3, S4, S5A). Consistent with its proposed landmark role, EXO70G1 localized to plasmodesmata in all genetic backgrounds examined, similar to its localization in wild-type plants (Figure 4A). Notably, PD localization of GFP-EXO70G1 in the *exo70A1* background suggests that its targeting does not require the canonical landmark subunit EXO70A1.

**Figure 4.**
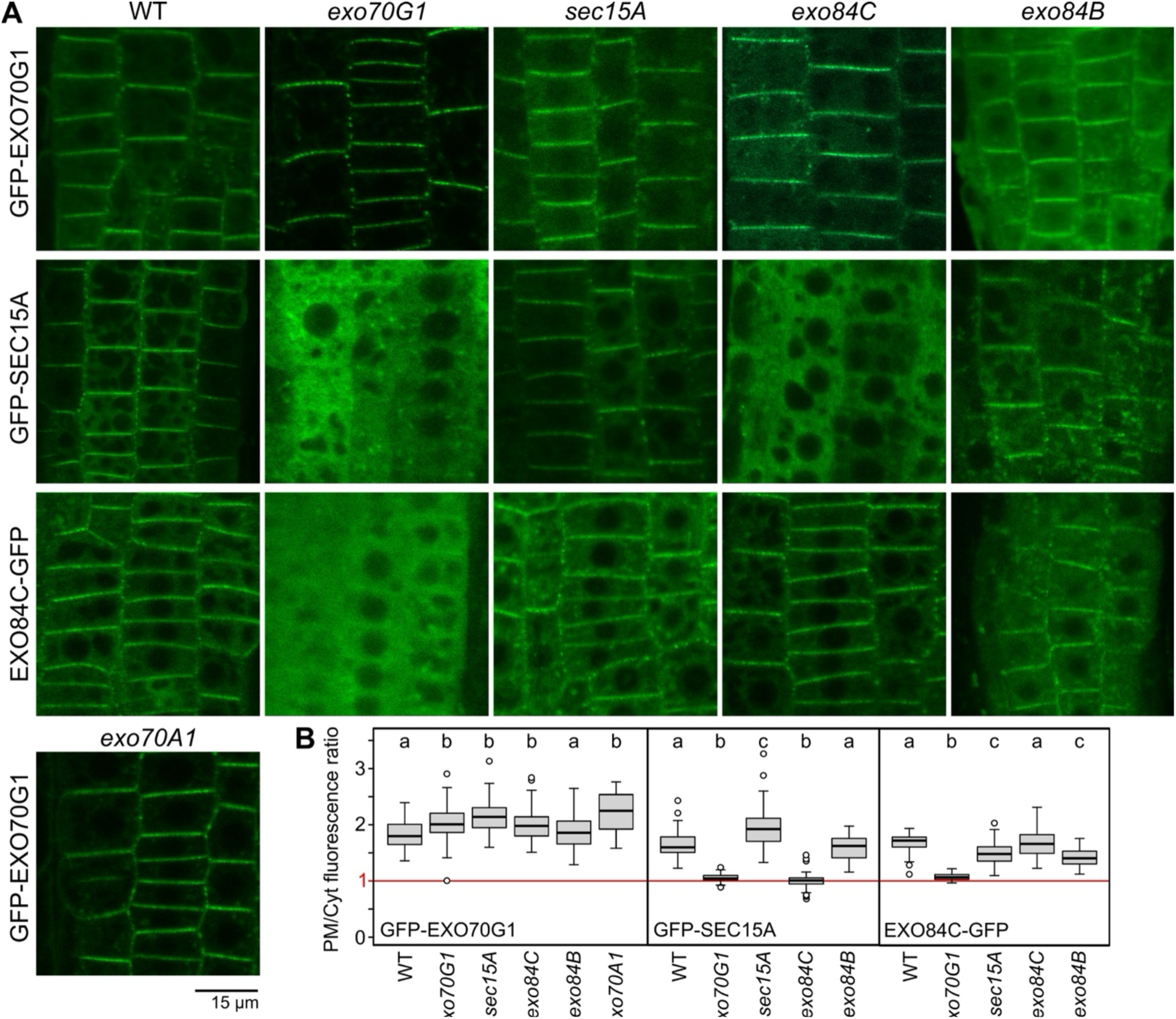
Effect of exocyst gene mutations on the localization of plasmodesmata-targeted subunits. **(A)** Localization of GFP-EXO70G1, GFP-SEC15A, and GFP-EXO84C in the root epidermis of 7-day-old wild-type seedlings and the indicated mutant backgrounds. **(B)** Plasma membrane–to–cytoplasm fluorescence ratio (PM/cyt ratio) for the indicated exocyst marker/host genotype combinations. At least 26 (typically >40) cells from at least four plants were analyzed per combination. Different letters indicate statistically significant differences (Kruskal–Wallis test, P < 0.05). All GFP-EXO70G1 transgenes were expressed under the control of the 35S promoter, and all other transgenes were expressed under their native promoters. A PM/Cyt ratio of 1 indicates a uniform distribution of GFP signal.

In contrast, loss of EXO70G1 strongly disrupted the PD localization of both SEC15A and EXO84C, redistributing them to the cytoplasm and intracellular membrane compartments. GFP-SEC15A was similarly mislocalized in the *exo84C* mutant (Figure 4A). Other mutant and transgene combinations had little, if any, effect on the localization of GFP-tagged exocyst subunits (Figure 4A). These qualitative observations were largely supported by the quantitative analysis of the plasma membrane–to–cytoplasm fluorescence ratio (Figure 4B).

We note that whereas EXO70G1 is expressed in all studied plant tissues, EXO70G2, a second EXO70G paralog in *Arabidopsis*, is barely expressed in most post-germination vegetative tissues under standard conditions (*26*, *27*). In our hands, GFP-tagged EXO70G2 could only be detected upon expression from the 35S promoter in the silencing-suppressed *rdr6* background (*28*), where it colocalized with callose at PD (Supplementary Figure S5B,C). Despite this, loss of EXO70G2, even in the *exo70G1 exo70G2* double mutant, did not result in obvious sporophytic phenotypes (Supplementary Figure S5A). We therefore excluded EXO70G2 from subsequent analyses of PD-exocyst localization and function.

Collectively, our results identify EXO70G1 as the principal determinant for recruitment of plasmodesmata-localized exocyst subunits.

### Lipid-dependent dynamic recruitment of the exocyst to plasmodesmata

Next, we analyzed molecular mechanisms of PD-exocyst recruitment to plasmodesmata. We first examined the dynamics of PD-associated GFP-tagged EXO70G1 and SEC15A using fluorescence recovery after photobleaching (FRAP). Both subunits exhibited rapid turnover, with a half-time of recovery of less than 3 s (Figure 5A), indicating continuous exchange between PD-associated and cytoplasmic pools rather than stable incorporation into a PD scaffold. This dynamic behavior suggested that steady-state localization of the exocyst at PD depends on ongoing membrane recruitment.

**Figure 5.**
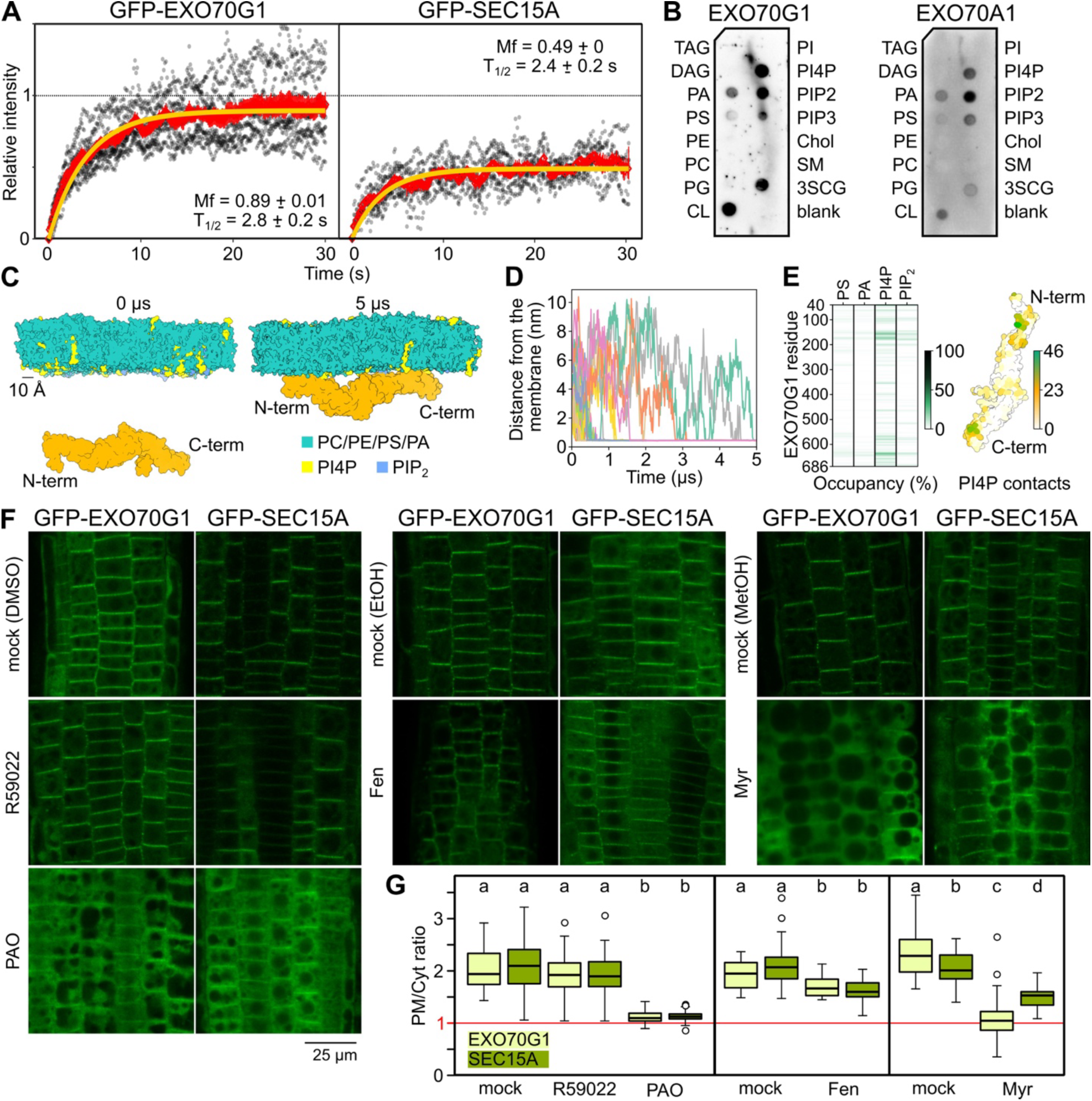
Plasmodesmata localization of exocyst subunits is dynamic and modulated by membrane lipid composition. **(A)** FRAP analysis of GFP-EXO70G1 and GFP-SEC15A at plasmodesmata in leaf epidermal cells. The half-time of fluorescence recovery is ∼3 s for both subunits. **(B)** Binding of *in vitro*–translated EXO70G1 and EXO70A1 to a panel of membrane lipids. TAG, triacylglycerol; DAG, diacylglycerol; PA, phosphatidic acid; PS, phosphatidylserine; PE, phosphatidylethanolamine; PC, phosphatidylcholine; PG, phosphatidylglycerol; CL, cardiolipin; PI, phosphatidylinositol; PI4P, phosphatidylinositol 4-phosphate; PIP_2_, phosphatidylinositol 4,5-bisphosphate; PIP_3_, phosphatidylinositol (3,4,5)-trisphosphate; Chol, cholesterol; SM, sphingomyelin; 3SCG, 3-sulfogalactosylceramide. **(C)** Representative snapshots of the molecular dynamics simulations of EXO70G1 and the phospholipid bilayer at the start of the simulation and the end of the simulation, with both the N-terminus and the C-terminus stably bound. **(D)** The progress of 20 independent MD runs, shown as plot of the minimal distance of EXO70G1 from the phospholipid bilayer. **(E)** A heatmap showing the percentage of time a lipid is found within the cutoff range of 8Å of each individual EXO70G1 residue (occupancy) and a depiction of PI4P occupancy as b-factor mapped onto the AlphaFold2 predicted structure of EXO70G1. **(F)** Effect of membrane lipid composition on the localization of fluorescent protein–tagged EXO70G1 and SEC15A in the root epidermis of 7-day-old seedlings. Images from inhibitor-treated plants are shown alongside solvent-treated controls. R59022, 12.5 µM, 1 h; PAO (phenylarsine oxide), 30 µM, 1 h; Fen (fenpropimorph), 50 µg mL⁻¹, 24 h; Myr (myriocin), 1 µM, 24 h. **(G)** Plasma membrane–to–cytoplasm fluorescence ratio for the indicated exocyst marker/treatment combinations. At least 20 (typically >50) cells from at least four plants were analyzed per condition. Different letters indicate statistically significant differences (one-way ANOVA, P < 0.05). A PM/cyt ratio of 1 indicates a uniform distribution of GFP signal. GFP-EXO70G1 was expressed under the 35S promoter, and GFP-SEC15A was expressed under its native promoter.

Because plasmodesmata are enriched in specific lipids, including sterols, sphingolipids, and signaling anionic phospholipids (*29–31*), we hypothesized that PD targeting of EXO70G1 may depend on its lipid-binding properties. To test this, we assessed the binding of *in vitro*–translated tagged EXO70G1 to a panel of membrane lipids using a protein–lipid overlay assay and compared it with canonical subunit EXO70A1. Both paralogs exhibited a similar, broad lipid-binding profile, with a strong preference for phosphoinositides and weak binding to phosphatidic acid (Figure 5B; Supplementary Figure S6).

To further investigate the membrane-binding mechanism of EXO70G1 with respect to its interaction with anionic lipids, we performed coarse-grained molecular dynamics (CG MD) simulations using the MARTINI2 force field (Figure 5C). Over 20 independent 5-μs simulations, EXO70G1 showed a clear preference for phosphatidylinositol 4-phosphate (PI4P) over phosphatidic acid (PA) or phosphatidylinositol 4,5-bisphosphate (PIP_2_; Figure 5D,E), in agreement with the protein-lipid overlay assay results. We next compared the membrane-binding mode of PD-associated EXO70G1 with the previously characterized plasma membrane-associated paralog EXO70A1 (*4*), which revealed both shared and distinct features of membrane interaction. In addition to the conserved C-terminal membrane interaction region shared by both EXO70G1 and EXO70A1, the simulations revealed an additional lipid-binding site in EXO70G1 near its N-terminus, enriched in positively charged arginine and lysine residues (Figure 5E).

To functionally assess the role of membrane composition in exocyst targeting to PD, we examined the effects of pharmacological perturbation of membrane lipid signaling or metabolism. Seedlings expressing GFP-tagged EXO70G1 or SEC15A were treated with: R59022 (a diacylglycerol kinase inhibitor affecting phosphatidic acid production), phenylarsine oxide (PAO; an inhibitor of phosphatidylinositol-4 kinases affecting PI4P production), fenpropimorph (an inhibitor of sterol biosynthesis), and myriocin (an inhibitor of sphingolipid biosynthesis). Consistent with our lipid-binding and computational simulation experiments, depletion of PI4P by PAO led to pronounced relocalization of both EXO70G1 and SEC15A from the plasmodesmata to the cytoplasm, while PA inhibition by R59022 had no effect (Figure 5F). Inhibition of sphingolipids by myriocin also led to a massive loss of membrane binding for both EXO70G1 and SEC15A. These effects were confirmed by quantification of the plasma membrane–to–cytoplasm fluorescence ratio, which additionally revealed partial delocalization of both subunits in fenpropimorph-treated plants (Figure 5G).

Together, these results indicate that the dynamic recruitment of EXO70G1 to plasmodesmata is supported by specific features of the PD membrane, particularly PI4P and sphingolipids, with sterols making a smaller contribution. The parallel relocalization of SEC15A upon lipid perturbation is consistent with EXO70G1 acting as a lipid-sensitive landmark for the recruitment of additional exocyst components to plasmodesmata.

### Exocyst at plasmodesmata controls callose turnover and symplastic transport

Targeted callose deposition and degradation are critical determinants of PD aperture size and depend on secretory and endomembrane trafficking pathways (*32*). We therefore asked whether disruption of PD-localized exocyst components affects plasmodesmatal callose accumulation and symplastic connectivity.

To test this, we first assessed symplastic connectivity in wild-type and *exo70G1* mutant seedlings by applying carboxyfluorescein diacetate (CFDA) to the cut cotyledon tips and monitoring fluorescence movement toward the root tip, as described in (*33*). Both visual inspection and quantification of CFDA fluorescence in the root meristem revealed reduced symplastic transport in the *exo70G1* mutant (Figure 6A). We corroborated this observation by analyzing callose deposition using aniline blue staining, which revealed a statistically significant accumulation of callose on rhizodermal and cortical cell walls in the root tips of *exo70G1* mutants (Figure 6B).

**Figure 6.**
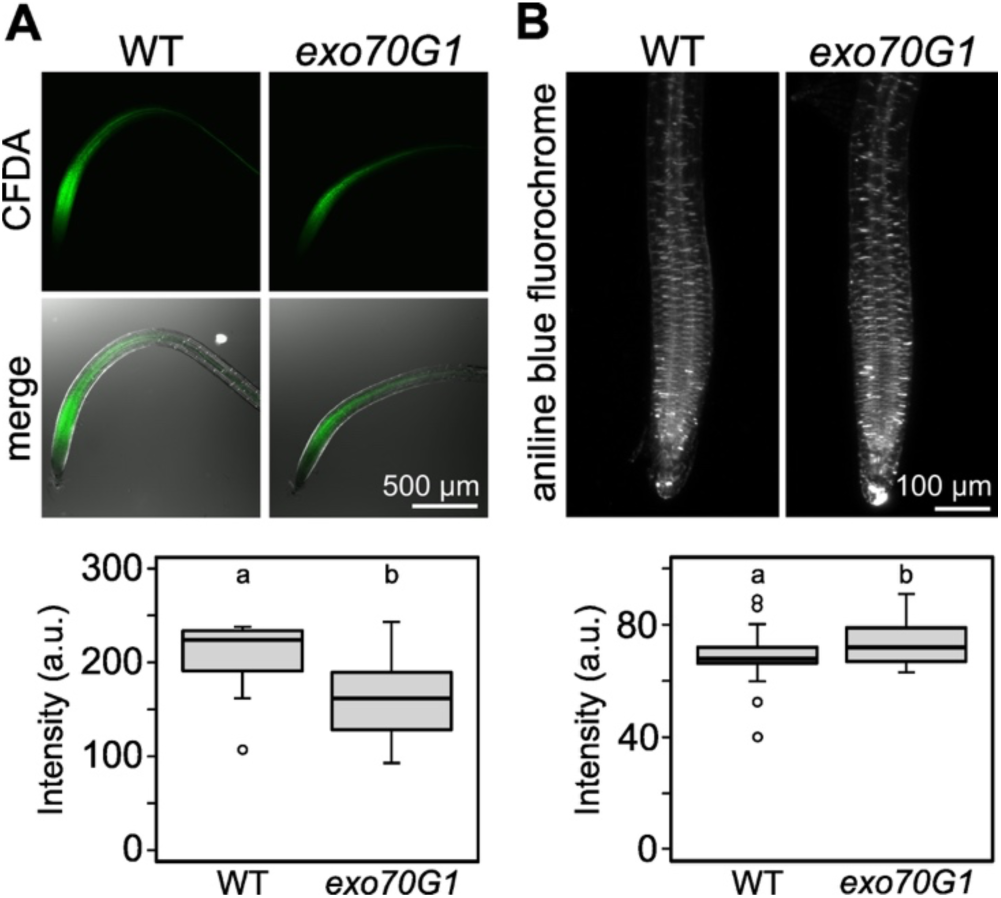
Perturbed symplastic connectivity and callose deposition in the *exo70G1* mutant. **(A)** Symplastic connectivity was assessed by applying 1 µL of CFDA solution to the cut cotyledon tip of 5-day-old wild-type (WT) and *exo70G1* seedlings. Fluorescence intensity in the root meristematic zone was quantified 2.5 h after application in roots from at least 14 plants per genotype. Different letters indicate statistically significant differences (Mann–Whitney test, P < 0.05). **(B)** Callose deposition in 7-day-old WT and *exo70G1* seedlings was visualized using aniline blue staining. Fluorescence intensity in the root meristem was quantified by averaging the signal from circular regions of interest (ROIs) of 50 µm diameter across 10 optical sections spanning the epidermal to cortical layers. At least 27 roots per genotype were analyzed. Different letters indicate statistically significant differences (unpaired t-test, P < 0.05).

To explore the functional consequences of callose overaccumulation in the absence of EXO70G1, we took advantage of the gain-of-function callose synthase mutant *cals3-3d*, which overaccumulates callose in the root stele, resulting in reduced root elongation and enhanced adventitious root formation, with only minor effects on shoot development (*34*). We generated a *cals3-3d exo70G1* double mutant and compared its phenotype with wild-type and single mutant plants at multiple developmental stages. In young seedlings, reduced root growth -already evident in *cals3-3d* - was strongly enhanced in the double mutant, whereas the *exo70G1* mutation alone had only a minor effect (Figure 7A).

**Figure 7.**
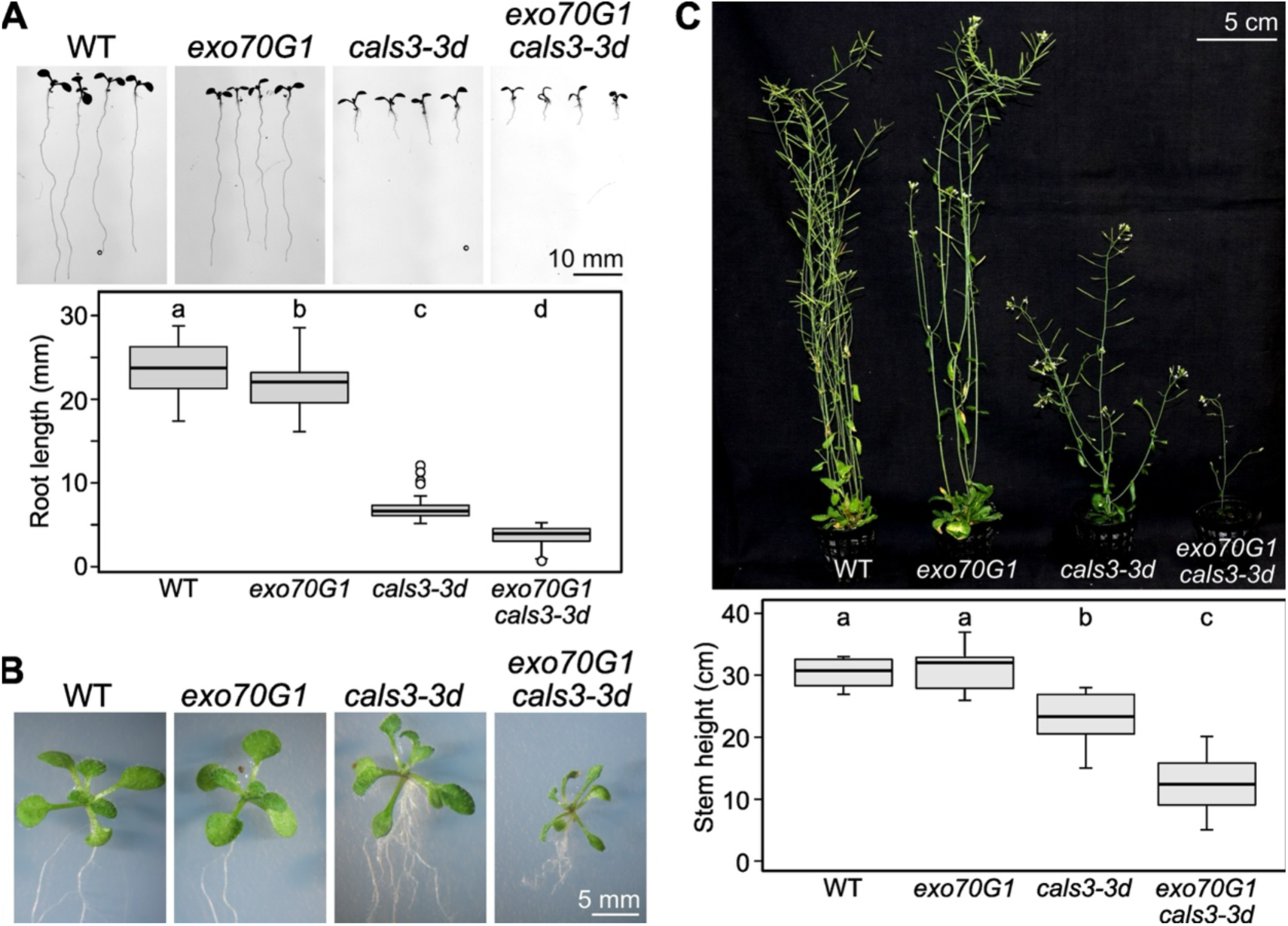
Loss of EXO70G1 aggravates the *cals3-3d* mutant phenotype. **(A)** Effects of *exo70G1*, *cals3-3d*, and the *cals3-3d exo70G1* double mutant on root length in 7-day-old seedlings. Root length was measured in at least 30 plants per genotype. Different letters indicate statistically significant differences (one-way ANOVA, P < 0.05). **(B)** Shoots of 14-day-old seedlings of the indicated genotypes. **(C)** Mature plants of the indicated genotypes. Shoot height was measured in at least 10 plants per genotype. Different letters indicate statistically significant differences (one-way ANOVA, P < 0.05).

In 2-week-old plantlets, the double mutant also exhibited markedly reduced shoot growth, likely reflecting impaired organ expansion (Figure 7B). Although double mutant plants initiated bolting, they developed into severely dwarfed and nearly sterile plants. In contrast, *exo70G1* single mutants appeared largely normal, with only a slight reduction in root growth compared to the WT plants, whereas *cals3-3d* single mutants were smaller and less fertile than WT plants and remained substantially less affected than the double mutants (Figure 7C). Similar phenotypes were observed for an independent *exo70G1* allele (*exo70G1-2*) in the *cals3-3d* background (Supplementary Figure S7).

Together, these observations indicate that loss of EXO70G1 (and hence the PD-localized exocyst) leads to reduced symplastic connectivity due to excessive callose accumulation at the plasmodesmata, which further aggravates the developmental defects of the gain-of-function *cals3-3d* mutant.

### Reduced plasmodesmal permeability in *exo70G1* mutants is associated with enhanced bacterial resistance

Regulated callose deposition at plasmodesmata is an important component of plant immune responses, as PD closure can limit the movement of pathogen-derived signals or effectors (*35*). Building on the observations above, we investigated the impact of *exo70G1* mutations on PD-dependent responses associated with plant defense.

We first examined short-distance cell-to-cell connectivity in leaves. Using particle bombardment, we introduced monomeric free GFP reporter into individual rosette leaf epidermal cells, while simultaneously marking transformed (source) cells by ER-targeted RFP expression. GFP movement into neighboring cells was then monitored in mock-treated plants and in plants treated with salicylic acid (SA), a key immune regulator known to suppress symplastic trafficking (*36*). The *exo70G1* mutant exhibited significantly reduced symplastic connectivity compared with the wild-type. SA treatment strongly inhibited symplastic transport in all genotypes examined (Figure 8A,B). Notably, SA also triggered relocalization of GFP-EXO70G1 and GFP-SEC15A from the PD to the cytoplasm, suggesting that the PD-associated exocyst pool itself responds to immune signaling (Figure 8C,D).

**Figure 8.**
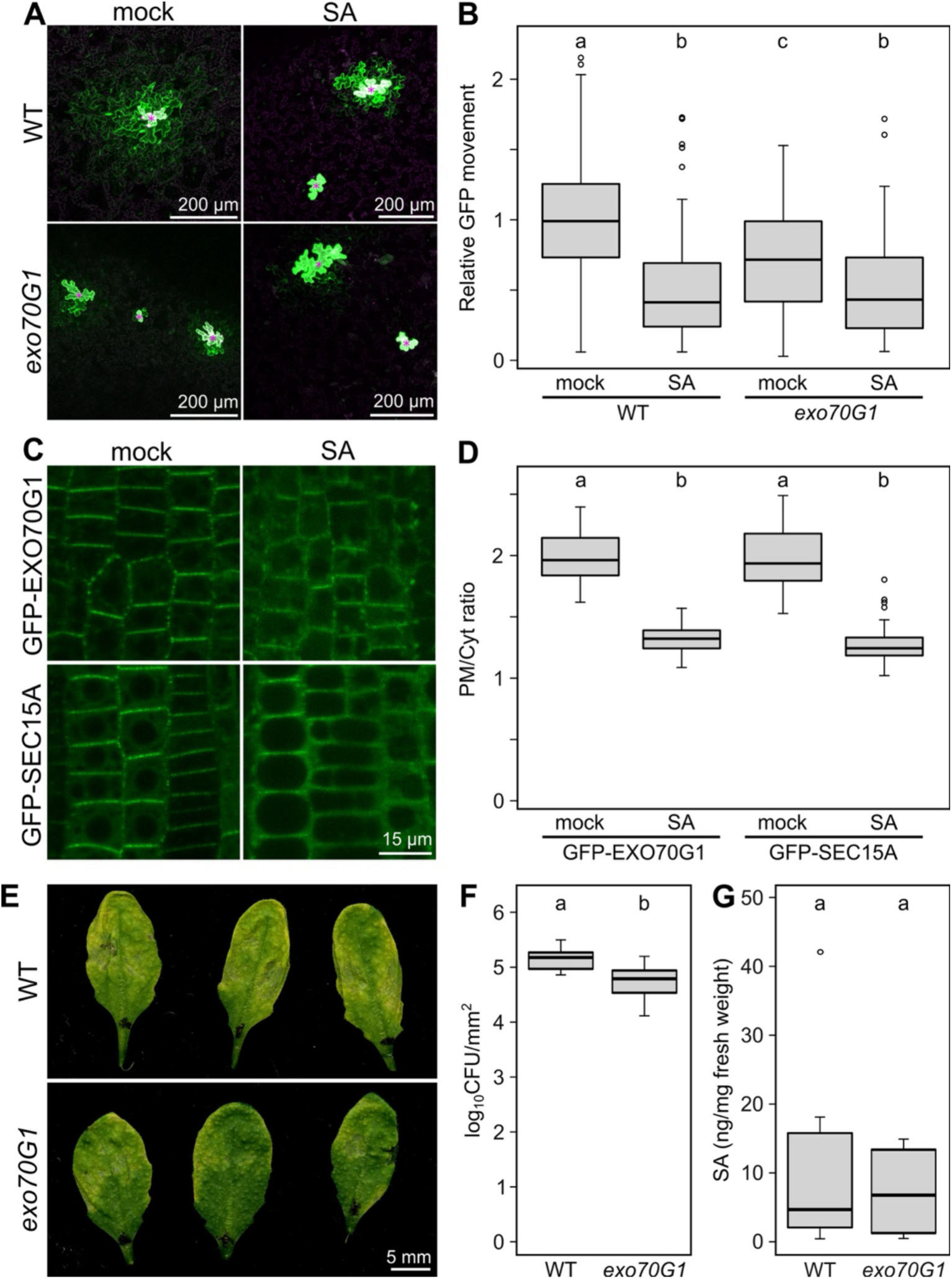
Defense-related phenotype of the *exo70G1* mutant. **(A)** Representative bombardment sites showing GFP movement from transformed cells (marked by purple asterisks) in mock-or SA-treated leaves of the indicated genotypes, 24 h after bombardment. Overlay of GFP (green) and RFP (magenta) channels is shown. **(B)** Quantification of relative GFP movement into neighboring cells in bombarded leaves of 4-week-old plants treated with water (mock) or 250 μM salicylic acid (SA) 1 h after bombardment. Data are cumulative from three biological replicates, comprising at least 180 (typically >200) individual bombardment sites per genotype and treatment. Different letters indicate statistically significant differences (one-way ANOVA, P < 0.05). **(C)** Effect of 250 μM SA on the localization of GFP–tagged EXO70G1 and SEC15A in the root epidermis of 7-day-old seedlings. Images from SA-treated plants are shown alongside mock-treated controls. **(D)** Plasma membrane–to–cytoplasm fluorescence ratio (PM/Cyt ratio) for the indicated exocyst marker/treatment combinations. At least 26 (typically >40) cells from at least four plants were analyzed per combination. Different letters indicate statistically significant differences (Kruskal–Wallis test, P < 0.05). **(E)** Disease symptoms two days after leaf infiltration with *Pseudomonas syringae* pv. *tomato*. **(F)** Quantification of bacterial load in infected leaves. Different letters indicate statistically significant differences (P < 0.05, unpaired t-test, n = 8). **(G)** Basal salicylic acid content in untreated leaves of the indicated genotypes. Differences between genotypes were not statistically significant (P > 0.05, unpaired t-test, n = 8).

Consistent with reduced symplastic connectivity, *exo70G1* mutant plants exhibited attenuated disease symptoms following inoculation with *Pseudomonas syringae* pv. *Tomato* (Figure 8E), accompanied by significantly reduced bacterial load (Figure 8F). These results indicate enhanced resistance of the mutant plants to the bacterial pathogen, which may be partially attributable to reduced PD permeability. Basal SA levels in the mutants were comparable to those in wild-type plants, ruling out constitutive SA accumulation as the cause for the observed enhanced resistance (Figure 8G).

Together, these results suggest that the PD-localized exocyst modulates symplastic connectivity in a manner relevant to bacterial infection, and that the enhanced resistance of *exo70G1* mutants is associated with reduced symplastic transport rather than constitutive activation of SA-dependent immunity.

### PD association is a feature of the EXO70G clade

Our data above demonstrated that EXO70G1, a member of the poorly characterized EXO70.3 subfamily, associates with plasmodesmata, whereas EXO70A1, belonging to the conserved EXO70.1 clade, does not. This raised the question of whether the PD association represents a unique characteristic of *Arabidopsis* EXO70G1 isoform or a more broadly conserved feature of related EXO70 homologs.

The exocyst is an ancient protein complex that predates the emergence of plasmodesmata. The EXO70.1 subfamily is considered closest to the ancestral EXO70 and it is functionally conserved (*10*), whereas EXO70.2 and EXO70.3 subfamilies form more dynamic and functionally divergent subfamilies (*12*, *13*). PD association is therefore unlikely to be an ancestral property but may have arisen within the EXO70.3 lineage during land plant diversification.

To assess the phylogenetic distribution of PD-associated EXO70 paralogs, we generated transgenic *Arabidopsis* lines expressing GFP-tagged heterologous EXO70 proteins representing distinct branches of the EXO70 family (Supplementary Figure S8). As a proxy for the ancestral, pre-multiplication state, we used the single EXO70 homolog from the filamentous alga *Klebsormidium nitens*. For the EXO70.3 clade, we selected the single EXO70 paralog from liverwort *Marchantia polymorpha*, as well as two paralogs from the ANA-grade angiosperm *Amborella trichopoda* (*37*), representing the EXO70G and EXO70I clades. The latter has been lost in Brassicaceae but retained in ferns, gymnosperms, and most other angiosperms (Supplementary Figure S8; (*10*, *38*).

When coexpressed with the PD-RFP marker, only *Arabidopsis* EXO70G1 and *Amborella* EXO70G displayed a characteristic punctate plasmodesmatal pattern and colocalized with the PD marker (Figure 9A,B). Quantitative PD index analysis further confirmed their enrichment at plasmodesmata (Figure 9C).

**Figure 9.**
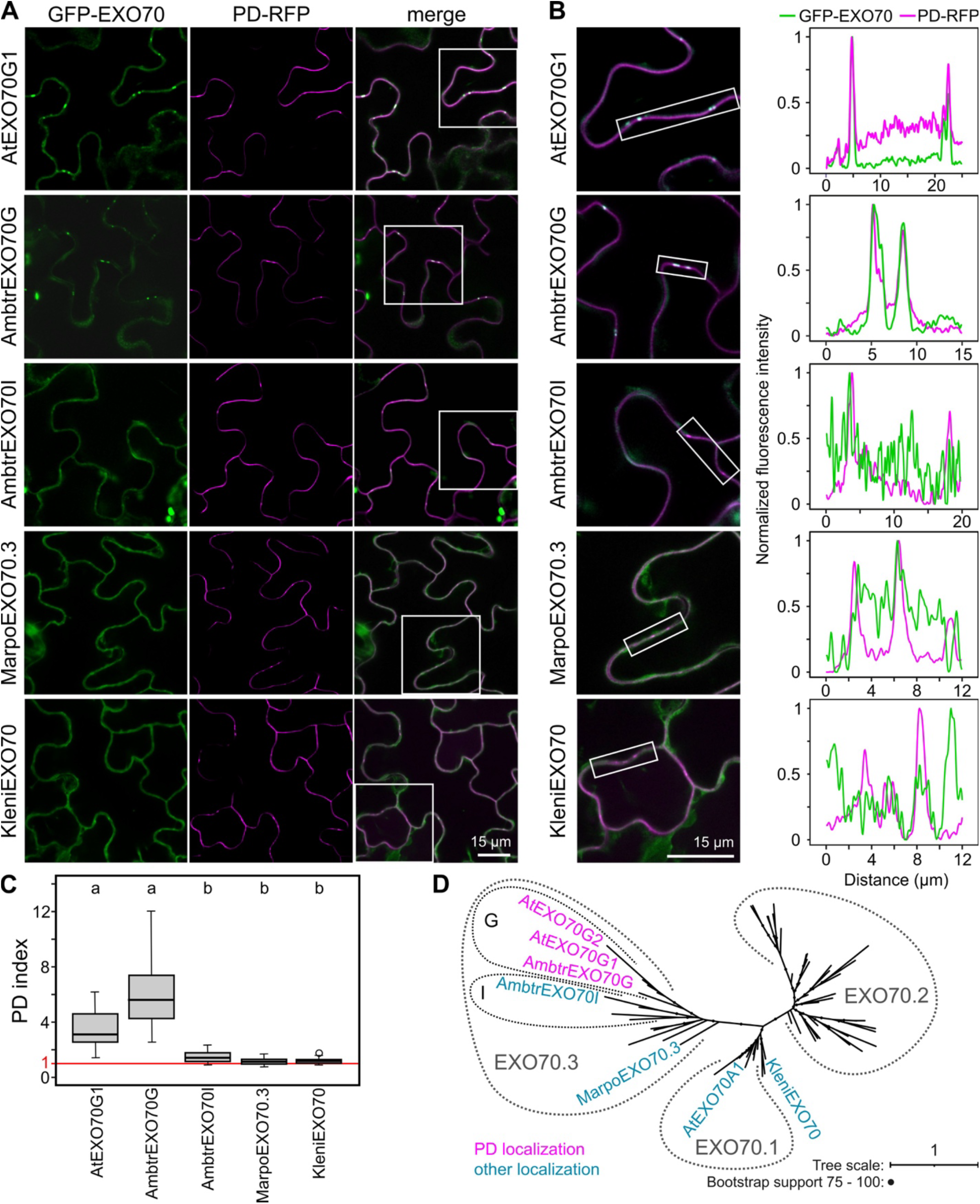
Phylogenetic distribution of plasmodesmata-localized EXO70 paralogs. **(A)** Localization of GFP-tagged heterologous EXO70 paralogs coexpressed with the PD-RFP marker in the cotyledon epidermis of 7-day-old *Arabidopsis thaliana* seedlings. At, *Arabidopsis thaliana*; Ambtr, *Amborella trichopoda*; Marpo, *Marchantia polymorpha*; Kleni, *Klebsormidium nitens*. All exocyst subunits were expressed under the control of the UBQ promoter, except for AtEXO70G1, which was expressed from the 35S promoter. Rectangles indicate regions shown in detail in **(B)**. **(B)** Magnified views of the regions indicated in **(A)**, shown together with intensity profiles of GFP and RFP fluorescence along the plasma membrane. **(C)** Plasmodesmata index (PD index) of the analyzed exocyst subunits, determined from at least 20 plasmodesmata per sample (four plants). Different letters indicate statistically significant differences (Kruskal-Wallis test, P < 0.05). **(D)** Schematic phylogenetic tree showing the distribution of plasmodesmata localization among EXO70 paralogs analyzed in this study (see Supplementary Figure S8 for the full tree).

These results indicate that PD association is a specific feature of the EXO70G clade consistent with the functional divergence of EXO70.3 subfamily into different clades in Euphyllophytes (Figure 9D).

## DISCUSSION

In this study, we identified a previously unrecognized form of exocyst module 2 that is associated with plasmodesmata and has a distinct subunit composition. We show that this specialized exocyst module contributes to the regulation of callose homeostasis, symplastic connectivity, and affect resistance to bacteria. Through comparative evolutionary analyses, we further demonstrate that its key landmark subunit EXO70G1 represents an evolutionarily derived plasmodesmata-targeting specialization within the EXO70 family in land plants. Our observations are consistent with previous proteomic analyses of plasmodesmata-enriched fractions, which detected several exocyst subunits. These include EXO70G1, SEC15A, and EXO84C (*39*); additionally, SEC15A was shown to form immobile plasma membrane puncta when expressed as a GFP fusion (*24*). Our *in vivo* colocalization and quantitative PD-index analyses now demonstrate conclusively that EXO70G1, SEC15A, EXO84C, and SEC10A specifically accumulate at plasmodesmata.

The subunit composition of such a PD exocyst module differs from the previously described canonical PM exocyst containing SEC15B, EXO84B, and EXO70A1 subunits, which function at the plasma membrane and during cell plate formation (*4*, *14*, *24*). Notably, our yeast two-hybrid and co-immunoprecipitation data further confirmed physical associations among the PD-associated exocyst subunits and indicated a preferential interaction between PD-exocyst subunit pairs, particularly EXO84C and SEC15A. While the presence of a complete hetero-octameric exocyst complex at PD remains unresolved, we speculate that PD-specific exocyst module 2 may associate with the evolutionarily less diversified and functionally generic module 1 at the PM/PD interface. This is corroborated by our observation that module 1 subunit SEC5A localizes to plasmodesmata and another module 1 subunit SEC3A interacts with PD-specific subunit EXO84C.

Analogously to the canonical PM exocyst pathway, in which EXO70A1 acts as a membrane landmark recruiting other exocyst subunits to the plant plasma membrane (*4*), EXO70G1 appears to perform a corresponding landmarking role in the PD exocyst formed by distinct module 2 subunits. Specifically, loss of EXO70G1 displaced SEC15A and EXO84C from PD, whereas EXO70G1 itself remained PD-localized in *sec15A* and *exo84C* mutants. The rapid FRAP recovery kinetics of both EXO70G1 and SEC15A at PD further indicate that this module is maintained by dynamic exchange rather than stable anchoring, consistent with regulatory flexibility. Interestingly, EXO70G1 also remains membrane-attached in the *exo70A1* mutant background, in sharp contrast to many other Arabidopsis EXO70 paralogs (*40*).

This difference may reflect a specific mode of EXO70G1–membrane interaction. Computational simulations and pharmacological perturbations suggested that EXO70G1 preferentially associates with PI4P over other anionic phospholipids and contains additional N-terminal lipid-binding sites not found in EXO70A1 (*4*, *41*). The qualitative *in vitro* protein–lipid overlay assays did not resolve this distinction, revealing broadly similar profiles for both proteins. In addition, EXO70G1–membrane interaction depends on sphingolipids and sterols, lipid classes enriched in plasmodesmata (*30*, *31*), whereas these lipids do not control EXO70A1 binding to the plasma membrane (*4*). Consistent with this lipid-dependent targeting model, an independent study reported that EXO70G1 accumulates preferentially at transverse plasma membrane domains in root epidermal cells, where it forms dynamic punctate foci with rapid exchange kinetics (*42*). The membrane association of EXO70G1 at these sites was sensitive to perturbation of sterol-dependent lipid nanodomains, suggesting that EXO70G1 targeting depends on local lipid composition rather than on conventional vesicular trafficking or cytoskeletal guidance. This lipid-dependent domain partitioning parallels the situation in trichomes, where EXO70A1 and EXO70H4 segregate to basal (PIP_2_-enriched) and apical (PA-enriched) domains, respectively, directing distinct secretory outputs (*21*). Collectively, these findings suggest that EXO70G1 relies on the evolutionarily conserved EXO70 lipid-recognition machinery, similarly to EXO70A1, but that its preferential accumulation at plasmodesmata reflects adaptation to the distinctive lipid composition of the PD membrane environment, rather than the emergence of a fundamentally different lipid-binding mechanism (see also (*43*) for other EXO70s).

The identity of EXO84C as a PD-exocyst subunit is also notable. Similarly to the division of labor between EXO70 isoforms described above, the EXO84B subunit serves as the major isoform in canonical PM secretory processes, including Casparian strip formation (*25*), whereas its paralog EXO84C has been previously linked to a VAP27-dependent pathway during stigma senescence (*44*). Its recruitment to PD, which depends on EXO70G1, suggests that EXO84C is a paralog associated with specialized membrane contact site functions rather than a redundant backup for EXO84B. Because plasmodesmata function is tightly linked to the ER–PM contact sites where MCTP protein tethers regulate molecular flux (*29*, *39*, *45*), the presence of EXO84C at PD may reflect a broader role for this isoform at unconventional membrane interfaces.

At the level of functional analyses, we found that the loss of EXO70G1 resulted in elevated callose accumulation and reduced symplastic transport, as demonstrated by both CFDA movement assays and particle bombardment GFP mobility assays. Moreover, *exo70G1* strongly enhanced the developmental and transport phenotypes of the callose-overproducing *cals3-3d* mutant (*34*), genetically linking EXO70G1 function to callose-dependent PD regulation. These observations strongly indicate that the EXO70G1-centered exocyst helps maintain PD in a conductive state by promoting callose turnover. Although the direct molecular cargoes of the PD-exocyst remain to be identified, it is tempting to speculate that its pathway acts through the trafficking of factors that control callose production, such as callose-degrading β-1,3-glucanases (*46*), negative regulators of callose synthase activity like CRK2 (*47*) or FW2.2 homologs (*48*), or other PD-associated components (*3*).

One surprising consequence of PD-exocyst-mediated callose regulation is evident in pathogen defense. Loss-of-function *exo70G1* mutants displayed enhanced resistance to *Pseudomonas syringae* despite normal basal salicylic acid levels, indicating that the resistance does not arise from constitutive immune activation. We speculate that increased callose deposition and reduced PD permeability may restrict the symplastic movement of susceptibility factors or signaling molecules required for efficient bacterial colonization. Alternatively, reduced PD aperture may itself be perceived as a stress signal, leading to defense priming. In this regard, our data provide a mechanistic, biologically relevant framework that corroborates the recent findings of (*35*), which show that plasmodesmal closure via synthetic tools activates stress signaling pathways and enhances plant resistance. The relocalization of EXO70G1 and SEC15A from PD upon SA treatment further indicates that the PD-associated exocyst is responsive to immune signaling. Together, these observations link PD-localized exocyst function to the broader regulation of plant immunity by controlling symplastic connectivity.

Finally, heterologous expression of EXO70 proteins from diverse land plant lineages revealed that PD targeting is a derived feature of the EXO70G clade within the EXO70.3 subfamily. Only *Arabidopsis* EXO70G1, EXO70G2 and *Amborella* EXO70G localized to the PD, whereas the *Klebsormidium* EXO70 (representing the pre-expansion ancestral state), *Marchantia* EXO70.3, and *Amborella* EXO70I did not. While these data were obtained in a heterologous system and should be interpreted with caution, they are consistent with our observations that *Marchantia* EXO70.3 localizes to the cytoplasm and plasma membrane in *Marchantia* tissues (*10*). The related EXO70I clade, absent from *Brassicaceae* and *Marchantia polymorpha* (*38*), appears to have followed a different specialization route, as *Medicago* EXO70I functions in periarbuscular membrane formation during mycorrhizal symbiosis (*49*, *50*). Thus, EXO70.3 paralogs diversified toward distinct specialized membrane interfaces (*10*, *13*), with PD targeting arising specifically within the EXO70G branch after its divergence from other EXO70.3 members.

In summary, our study links secretory pathway regulation to callose homeostasis, symplastic connectivity, plant development, and defense. We further show that this regulatory mechanism is associated with the evolutionary diversification of plant exocyst subunits. Whereas previous studies of plant exocyst specialization have focused primarily on functional diversification among EXO70 paralogs, our data indicate that the PD-associated exocyst variant comprises multiple specialized subunits, including EXO70G1, SEC15A, and EXO84C. Together, these subunits define a distinct exocyst module 2 adapted to the plasmodesmal membrane environment. These findings corroborate and extend our previous model (*51*), showing that plants diversified not only individual EXO70 landmarks but also broader exocyst assemblies adapted to distinct membrane environments and specific cellular interfaces. Identifying the cargoes and regulatory protein partners that connect EXO70G1-dependent vesicle tethering to callose turnover at plasmodesmata represents an important objective for future investigation.

## MATERIALS AND METHODS

### Arabidopsis thaliana *lines*

All lines used were derived from the *A. thaliana* Columbia-0 (Col-0) ecotype. The following T-DNA insertion alleles were used for *EXO70G1* (At4g31540): *exo70G1-1* (SALK_099892, this line is referred as *exo70G1* throughout the study), obtained from the SALK institute collection (*52*), *exo70G1-2* (GK_209C09) for EXO70G2 (At1g51640): *exo70G2* (GABI_548B11) from the GABI-KAT collection (*53*). Some experiments involved the previously described *exo84C-1* (*25*), *sec15A* (*24*), *exo70A1-1* (*54*), and *exo84B-1* (*14*) mutants. The presence of T-DNA insertions in these plants and the progeny of their crosses was verified by PCR (Supplementary Figure S3) using the primers listed in Supplementary Table S1. Absence of the EXO70G1 protein in homozygous *exo70G1-1* plants was verified by immunoblot analysis (Supplementary Figure S3C). In addition, plants carrying the chemically induced *cals3-3d* mutation (*34*), the *rdr6* mutation (*28*), and the PD-marker, mRFP tagged At5g24010 (*22*) were employed in some experiments.

### Cloning and transgenic plant construction

All PCRs were performed using Q5 polymerase (New England Biolabs). Constructs for yeast two-hybrid assay were amplified from *Arabidopsis* Col-0 cDNA. Regions of AtEXO84C were amplified using specific primers flanked by XmaI (full-length and EXO84C-C) or EcoRI (EXO84C-N) and SalI sites. Amplified products were introduced into vectors pGADT7 and pGBKT7 (Clontech) to obtain AD- and BD-fused AtEXO84C, respectively.

To create GFP-tagged AtEXO84C, DNA fragments were amplified from *Arabidopsis* Col-0 genomic DNA using specific primers and cloned into pENTR3C vector (Invitrogen) via KpnI and NotI restriction sites. Entry clone was recombined into the Gateway binary vector pH7FWG,0 (*55*) to generate pAtEXO84C::AtEXO84C:GFP.

For cloning of the GFP-tagged AtEXO70G1 and AtEXO70G2, the MultiSite Gateway approach was used. The AtEXO70G1 promoter (2779 bp upstream of start codon) and AtEXO70G1 and AtEXO70G2 coding sequences (CDS) were amplified using specific primers from *Arabidopsis* Col-0 genomic DNA and subcloned by BP Clonase (Invitrogen) into pDONR P4-P1r and pDONR P2r-P3 (Invitrogen), respectively. To produce the final construct pAtEXO70G1::GFP:AtEXO70G1, entry clones containing AtEXO70G1 promoter and AtEXO70G1 CDS were assembled together with pEN-L1-F-L2 (*56*) and destination vector pB7m34GW (*57*) by LR Clonase II Plus (Invitrogen). To produce 35S::GFP:AtEXO70G1 and 35S::GFP:AtEXO70G2, entry clone containing 35S promoter pEN-L4-2-R1 (*56*), AtEXO70G1 or AtEXO70G2 CDS, pEN-L1-F-L2 and pB7m34GW were used. CDS of *Amborella trichopoda* EXO70G and EXO70I were synthesized (Eurofins Genomics) and cloned into pAtUBQ10:eGFP-MCS using BamHI and PacI restriction sites.

Constructs for *in vitro* coupled transcription and translation reaction (HA-AtEXO70A1 and HA-AtEXO70G1) were prepared using specific primers according to (*58*).

Transgenic *Arabidopsis thaliana* plants expressing tagged proteins were generated using the floral dip method (*59*). Combined genotypes were obtained by crossing the indicated mutant and transgenic lines.

### Culture conditions

For *in vitro* cultures, *Arabidopsis* seeds were surface-sterilized for 10 min in 20% household bleach (Bochemie, Savo), rinsed three times with sterile distilled water, stratified for 2-4 d at 4°C and germinated on vertical plates containing ½ Murashige and Skoog salts (Sigma-Aldrich) supplemented with 1% (w/v) sucrose (Fluka/Sigma-Aldrich), vitamins, and 1.6% (w/v) plant agar (all Duchefa), buffered to pH 5.7. Seedlings were grown vertically in a climate chamber at 22 °C under long day (16 h light, 8 h dark) conditions. Depending on the assay, 5-, 7- or 10-day-old seedlings were used. Where indicated, young seedlings were transferred to Jiffy pellets and moved to *ex vitro* (also 22 °C, under long day conditions) for further phenotypic studies, crossing, floral dip method or seed amplification.

For leaf symplastic connectivity measurements, bacterial inoculation assay, and salicylic acid contents analyses, plants were cultivated under short day conditions (10 h light,14 h dark), at 22 °C and 70 % humidity.

### Aniline blue staining

Callose was stained by incubating whole seedlings in aniline blue fluorochrome solution (Biosupplies), diluted 1:3 in ½ MS medium, for 1.5 h. Confocal images were acquired using a 10× objective. Fluorescence intensity was quantified by averaging signal from circular regions of interest (ROIs) of 50 µm diameter across 10 optical sections spanning the epidermal to cortical layers in the root meristem. At least 27 roots per genotype were analyzed.

For colocalization of GFP-tagged proteins with aniline blue, 10-day-old seedlings were mounted in aniline blue solution (as described above) and imaged using a Mirava Polyscope (Abberior).

### Microscopy and image analysis

Subcellular localizations of fluorescently tagged proteins were examined in young *Arabidopsis* roots (7d), cotyledon (7d) or leaves (10d) using ZEISS LSM 900 (JENA/Germany) equipped with Airyscan 2 and using a 488/561 nm laser (objective: LD LCI Plan-Apo 40x/1,2 Imm Corr DIC M27, Plan-Apochromat 63x/1.40 Oil DIC M27).

The *exo70G1* seedlings crossed with *cals3-3d* were imaged on Stereomicroscope Leica M205FA (objective Plan-Apochromat 2x, camera Leica DMC6200 colour 2.2Mp)

Images of GFP tagged exocyst subunits with aniline blue fluorochrome colocalization were taken on Mirava Polyscope (Abberior) using 60× water-immersion objective (NA 1.20, UPLXAPO60XW). eGFP channel: Fluorophores were excited with a 488 nm laser at 15% power and detected using an APD detector with a 498–647 nm detection window. Time-gated detection was used (1.531 ns + 7.219 ns). Aniline Blue 405 channel: Excitation was performed with a 405 nm laser at 15% power, detected using an APD detector with a 415–478 nm detection window.

Post-acquisition image processing and quantification were performed using Fiji software (for GFP and CFDA quantification) (*60*) or ZEN Blue (Zeiss) (for aniline blue fluorochrome quantification). Figures were prepared in Inkscape. The PM association was calculated as a ratio of the PM/cytoplasm GFP signal based on mean gray values in narrow regions of the PM and next to it in the cytoplasm, avoiding vacuoles or nuclei. PD index was calculated as a ratio of the GFP signal in PD/GFP signal on the PM next to the PD based on mean gray values. The region of PD was selected according the mRFP tagged PD marker.

Fluorescence intensity profiles of different fluorescently tagged proteins along the plasma membrane were plotted in R (4.5.2) using the ggplot2 package (4.0.0). Intensity was normalized independently for each channel using the 5th and 99th percentile intensities along the profiles as lower and upper reference values, respectively, to avoid the influence of outliers. Intensities were rescaled to the [0,1] range.

Fluorescence recovery after photobleaching (FRAP) was performed using a confocal microscope ZEISS LSM 900 (JENA/Germany). A defined region of interest at plasmodesmata was photobleached using high laser power, and fluorescence recovery was monitored at 0.12 s intervals. Data were normalized to pre-bleach intensity and used to calculate the half-time of recovery (t½). 10 cells from 3 plants were analyzed for each genotype. The experiment was repeated 2 times with similar results.

### Yeast 2-hybrid system

For the Y2H assay we employed the Matchmaker GAL4 Two-Hybrid System 3 (TaKaRa) and followed the manufacturer′s protocol. Some constructs carrying genes encoding for exocyst subunits were prepared previously (*24*, *61*, *62*). Cells of *Saccharomyces cerevisiae* AH109 were transformed with pairs of plasmids for tested exocyst subunits or controls (empty plasmids) and selected on -LEU -TRP medium. Single colonies were then scale diluted in distilled water and grown for 3 d at 28°C on -ADE -HIS -LEU -TRP selective medium. At least two repetitions were made. The EXO84C-N fusion to the DNA-binding domain resulted in autoactivation activity. Consequently, only activation domain fusions were used for the full-length EXO84C, while DNA-binding domain constructs of the N- and C-terminal halves of EXO84C were generated and employed in this assay. DNA-binding domain fusions to SEC3 and SEC10B were completely omitted due to their autoactivation activity.

### Protein co-immunoprecipitation analysis and mass spectrometry analysis

In vivo interactions of exocyst subunits were detected by co-immunoprecipitation using the µMACS GFP-Tagged Protein Isolation Kit (Miltenyi Biotec) according to the manufacturer’s instructions, with slight modifications as described below. One gram of 10-day-old *Arabidopsis* seedlings expressing GFP-tagged bait subunits was ground in liquid nitrogen, and 1 mL of Sec6/8 buffer was added (20 mM HEPES, pH 6.8, 150 mM NaCl, 1 mM EDTA, 1 mM DTT, 0.5% Tween-20, 1:100 protease inhibitor cocktail; Sigma-Aldrich). The Sec6/8 buffer was used instead of the provided lysis buffer. The lysate was centrifuged at 10,000 × g for 10 min at 4°C. Anti-GFP magnetic beads (100 μL) were added to the supernatant and incubated on ice for 30 min. The solution was then transferred to a column attached to a magnetic stand and washed four times with 200 μL of Sec6/8 buffer, followed by one wash with 100 μL of washing buffer 2. Bound proteins were eluted with 100 μL of preheated elution buffer, frozen in liquid nitrogen, and stored at −80°C. Eluted proteins were analyzed by liquid chromatography coupled to tandem mass spectrometry (LC-MS/MS). For a detailed description of the LC-MS/MS analysis, see (*4*). Results were processed using Data Analysis 4.1 and the Mascot server to obtain Normalized Spectral Abundance Factor (NSAF) values reflecting the specificity of the detected protein interactions.

### Protein–lipid overlay assays

To test whether EXO70G1 and EXO70A1 show preferential binding for some lipids, the specific EXO70G1 and EXO70A1 constructs were generated and used according to the manufacturer’s instructions as templates for *in vitro* coupled transcription/translation using the TNT SP6 High-Yield Wheat Germ Protein Expression System (Promega) in a total volume of 50 ul. The reaction product N-terminal HA-tagged EXO70G1 or EXO70A1 (Supplementary Figure S6) were used for lipid strips (Echelon Biosciences). Membrane lipid strips (Echelon Bioscience, P-6002) were blocked with 3 mL blocking solution (3% bovine serum albumin [Sigma, A7030], 0.1% Tween-20 in phosphate-buffered saline [PBS], pH 7.2) under gentle shaking for 30 min at RT. Then, HA-tagged EXO70G1 or EXO70A1 were added (final protein concentration of 2 µg/mL) to lipid strip with blocking solution and incubated for 1.5 h. Strips were then washed three times for 10 min with washing solution (PBS containing 0.1% Tween-20). Anti-HA antibody (Invitrogen, 26183) diluted 1:1000 in the blocking solution was applied on strips and incubated at RT for 1 h under gentle shaking, followed by three washes and subsequent 50 minutes incubation with secondary antibody diluted 1:10000 (Promega, W4021). After three washes for 10 min with washing solution, strips were incubated with 600 µL Amersham ECL Prime Western Blotting Detection Reagent (GE Healthcare, RPN2236) for 2 min, and bound proteins were visualized using the ChemiDoc XRS+ Imaging system (Bio-Rad).

### Molecular dynamics simulations

To examine the membrane binding mode of EXO70G1, we performed molecular dynamics (MD) simulations of EXO70G1 and a phospholipid bilayer using the MARTINI2 force field (*63*, *64*). A coarse-grained bilayer, composed of POPC:POPE:POPS:POPA:POPI4P:POPI(4,5)P2 (ratio 37:37:10:10:5:1) as previously described in (*4*), was generated using the Python insane.py script (*65*). An AlphaFold2 model of EXO70G1 was accessed via the AlphaFold Protein Structure Database (*66*, *67*), manually trimmed to exclude structures with low confidence and converted into a coarse-grained representation using the martinize2.py script (*68*) with the Elastic Network in Dynamics (ElNeDyn) extension to preserve secondary structure (*69*). All simulations were performed using GROMACS (version 2021.4; (*70*)). Detailed MD parameters are included in (Supplementary Data S2 – a table with MD parameters). Protein and lipid contacts were analyzed using the PyLipid Python package (*71*) and visualized with custom Python scripts.

### Western blot analysis

Western blot analysis was performed according to standard procedures, using the primary antibodies anti-EXO70G1 mouse (polyclonal, dilution 1:1000), anti-GFP mouse (AS152987, Agrisera, dilution 1:1000) and Horseradish peroxidase-conjugated secondary antibodies (anti-mouse 1:10000; Promega). After the washing step the membrane was incubated with 1000 µL Amersham ECL Prime Western Blotting Detection Reagent (GE Healthcare, RPN2236) for 2 min and visualized using the ChemiDoc XRS+ Imaging system (Bio-Rad).

### Symplastic transport and connectivity assays

Cell-to-cell connectivity was assessed by microprojectile bombardment of pB7WG2.0.RFPER and pB7WG2.0.GFP plasmids into the abaxial side of fully expanded leaves of 4-week-old plants, as described in (*72*). One hour after bombardment, leaves were infiltrated with salicylic acid (SA 250 µM), or dH_2_O (mock). Leaves were kept in Petri plates with MS1/2 medium containing 0.7% agar at room temperature overnight before imaging. RFP was excited with a 561-nm DPSS laser and collected at 600 to 640 nm, while GFP was excited with a 488-nm argon laser and collected at 505 to 530 nm using a Plan-Apochromat 10x/0.45 M27 objective. The number of GFP-positive cells was counted manually, with each bombardment site considered as independent value; 3-10 bombardment sites were assessed per leaf, 3 leaves per plant, 3 plants per treatment.

### Pseudomonas inoculation assays

*Pseudomonas syringae pv. tomato* DC3000 (*Pst*) was grown overnight on plates containing LB medium (tryptone 10 g/L, NaCl 10 g/L, yeast extract 5 g/L, pH 7.0) supplemented with 1.4% agar and 50 mg/L rifampicin. Three similarly sized mature leaves from four-week-old plants (typically the 8^th^ to 10^th^ leaf) were syringe-infiltrated with a suspension of bacteria (OD_600_=0.05 in 10 mM MgCl_2_). Two days after infection, one 6 mm diameter disc from each treated leaf was cut using a corkborer; three leaf discs from one plant were pooled into one sample and homogenized in 2 mL Eppendorf tubes with 1 mL 10 mM MgCl_2_ and 1 g of silica beads using a FastPrep-24 instrument (MP Biomedicals, USA). The resulting homogenate was subjected to serial 10× dilutions and pipetted onto LB plates. The colonies were counted after 2 days of incubation at 26°C. Eight individual plants were used per treatment, bacterial load being expressed as log10(CFU*mm^−2^) (*73*).

### Salicylic acid (SA) quantification

Free SA was quantified using a modification of the *Acinetobacter* sp. ADPWH_lux bioassay (*74*). Bacteria were grown in liquid LB medium without antibiotics overnight at 37 °C on a rotary shaker (120 rpm). At the day of measurement, fresh LB medium was mixed with bacterial culture (approx. 1:10) and cultivated at the same conditions until OD_600_=0.4.

Plant material was collected into 2 mL Eppendorf tubes with 1 g of 1.3 mm ceramic beads and frozen in liquid nitrogen (50-150 mg FW). For the calibration curve preparation, 6 extra samples of untreated WT plants were harvested. Frozen samples were homogenized in FastPrep-24 instrument (MP Biomedicals, USA) (40 s, 4.5 m/s) and re-frozen in liquid nitrogen. Then, NaAC (0.1 M, pH 5.6) buffer was added (250 µL per 100 mg FW) and the homogenization step was repeated. Samples were centrifuged for 10 min at 15000 g at 4°C, supernatant was transferred into fresh tubes and kept on ice. Next, in sterile white microtiter plate with opaque bottom were mixed 60 µl of LB, 50 µl of *Acinetobacter* sp. ADPWH_lux culture (OD_600_=0.4) and 20 µl of plant extract or standard, with three technical replicates. Mixture was incubated 1 hour at 37 °C and luminescence was read on Tecan Infinite microplate reader (Austria), well integration time 4 s. Standards were prepared by adding the known amount of SA (range 1-50 ng/mL) into untreated WT plant extract. SA content was quantified based on a linear fitted calibration curve. Samples were assessed in 4 technical and 8 biological replicates.

### Phylogenetic tree construction

The EXO70 protein sequences were identified as described previously (*10*). An initial alignment has been produced by MAFFT L-INS-i (*75*) and manually trimmed. A maximum likelihood phylogenetic tree was constructed using Seaview (*76*) and validated using the bootstrap method with 500 replicates.

### Statistics

All experiments were repeated at least three times (data from a representative replication are shown). Statistical analyses were performed using GraphPad Prism 8, the RStudio environment (cell-to-cell connectivity), or an online calculator (*77*). The P values were calculated using a two-tailed Student’s t-test, one-way or two-way ANOVA followed by the post-hoc Tukey HSD (honestly significant difference) test, or, alternatively, median Bootstrap, as stated in figure legends. Kruskal-Wallis test followed by the Dunn’s test with Holm correction or Mann–Whitney test were used to analyse non-normal distributed data.

## Acknowledgments

We would like to thank Jana Kaňerová for excellent technical assistance. We also express our gratitude to the developers of the open-source programs used extensively in this study, particularly FIJI, Gimp, GROMACS, Inkscape, R, Rstudio, VMD and UCSF ChimeraX.

## Author contributions

Conceptualization: E.J.D., V.Ž., M.P.

Investigation: E.J.D., S.H., T.K., M.V., E.Š, M.D., J.G.G., T.P., A.A., A.Z.

Resources: P.P., J.O., I.K., J.Š., R.P.

Formal analysis: E.J.D., K.J., F.C., M.P.

Supervision: E.J.D., M.P.

Funding acquisition: M.P.

Writing—original draft: E.J.D., F.C., M.P.

Writing—review & editing: all authors

## Data availability

All data needed to evaluate the conclusions in the paper are present in the paper and/or the Supplementary Data. Data files related to sequence alignment and phylogenetic tree reconstruction are deposited in the Zenodo repository (https://doi.org/10.5281/zenodo.21130885). The mass spectrometry proteomics data have been deposited to the ProteomeXchange Consortium via the PRIDE partner repository with the dataset identifier PXD080396.

## Supplementary Materials

Supplementary Figures S1 to S8

Supplementary Table S1

