## Supplementary Figures S1-S8 and Supplementary Table S1 for "A plasmodesmata-specific exocyst complex regulates symplastic connectivity by affecting callose turnover"

Edita Janková Drdová *et al.*

**This PDF file includes:**

Supplementary Figures S1 to S8
Supplementary Table S1

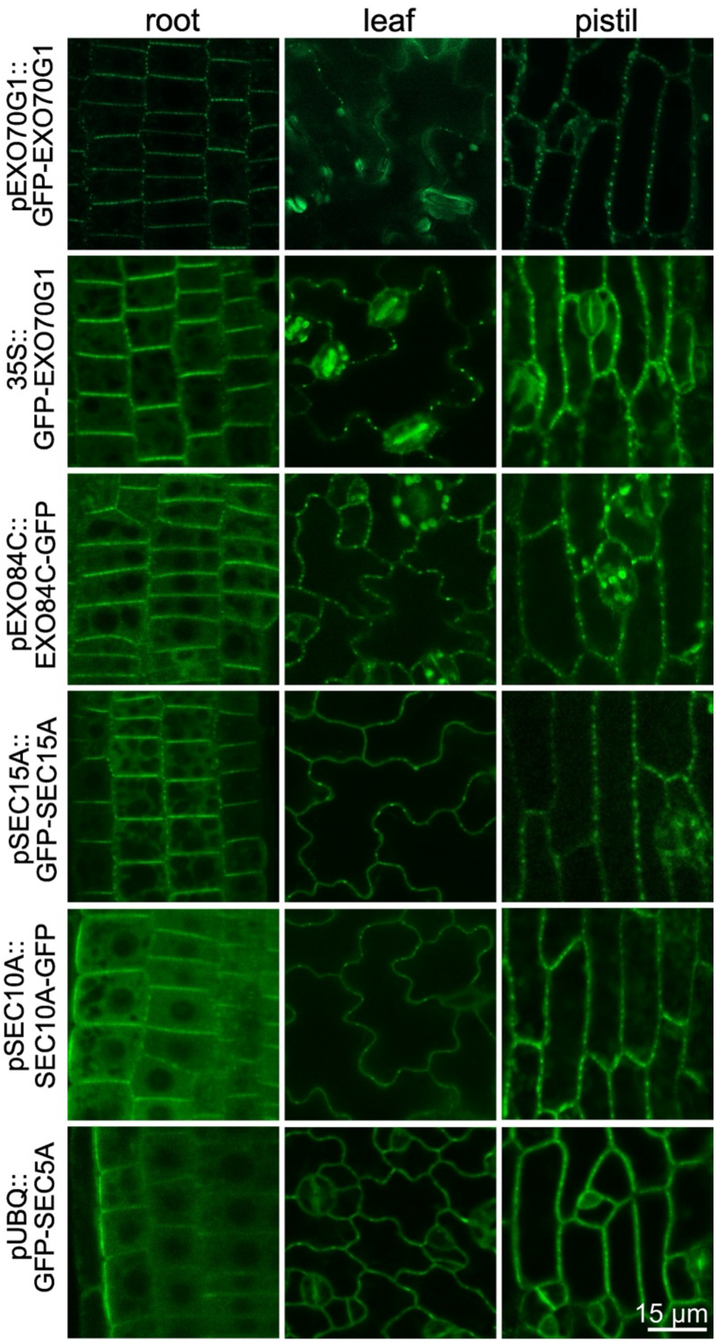


**Supplementary Figure S1.**

**Discontinuous plasma membrane localization of selected exocyst subunits suggests their association with plasmodesmata.** Localization of GFP-tagged exocyst subunits EXO70G1, SEC15A, EXO84C, SEC10A, and SEC5A in the indicated tissues of 7-day-old transgenic *Arabidopsis thaliana* seedlings. GFP-EXO70G1, EXO84C-GFP, GFP-SEC15A and SEC10A-GFP exocyst subunits were expressed under their native promoters; GFP-SEC5A was expressed under the control of the UBQ promoter, and GFP-EXO70G1 was also expressed under the 35S promoter.

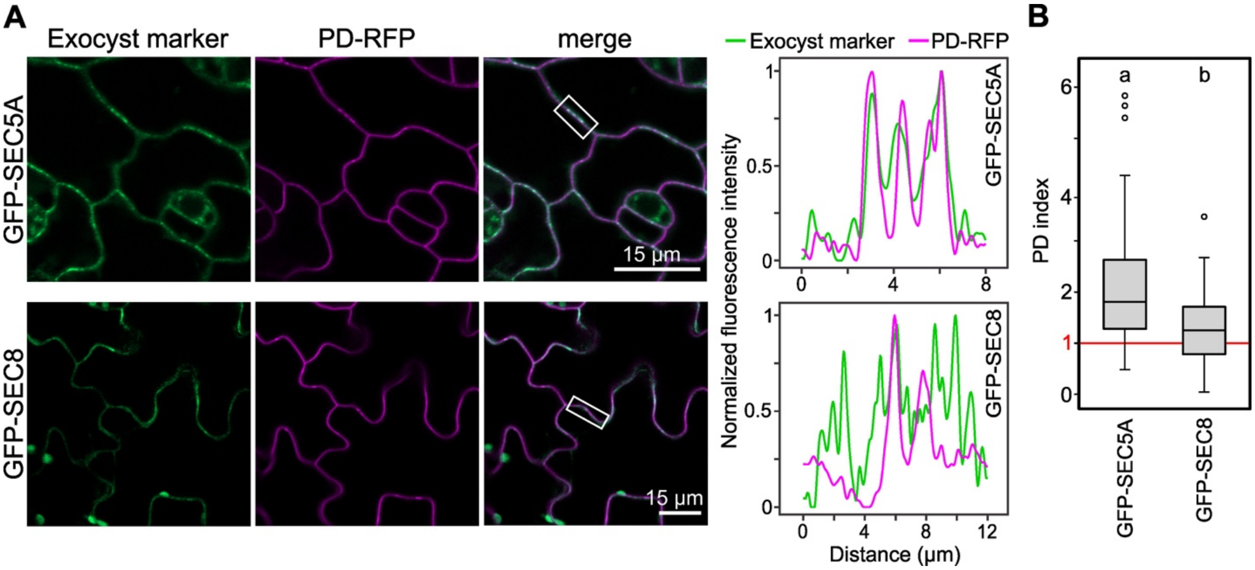


**Supplementary Figure S2.**

**SEC5A, but not SEC8, colocalizes with a plasmodesmata marker. (A)** Colocalization of GFP-tagged SEC5A and SEC8 with the PD-RFP marker in leaf epidermal cells of 10-day-old transgenic *Arabidopsis thaliana* seedlings. Intensity profiles of GFP and mRFP fluorescence along the plasma membrane are shown for the region indicated by the rectangle. **(B)** Plasmodesmata index (PD index) was determined from at least 28 plasmodesmata per sample (four plants). Differences between genotypes were statistically significant (Mann-Whitney test, $P < 0.01$). A PD index value of 1 indicates a uniform distribution of GFP signal along the plasma membrane. SEC5A was expressed under the control of the UBQ promoter, whereas SEC8 was expressed under its native promoter.

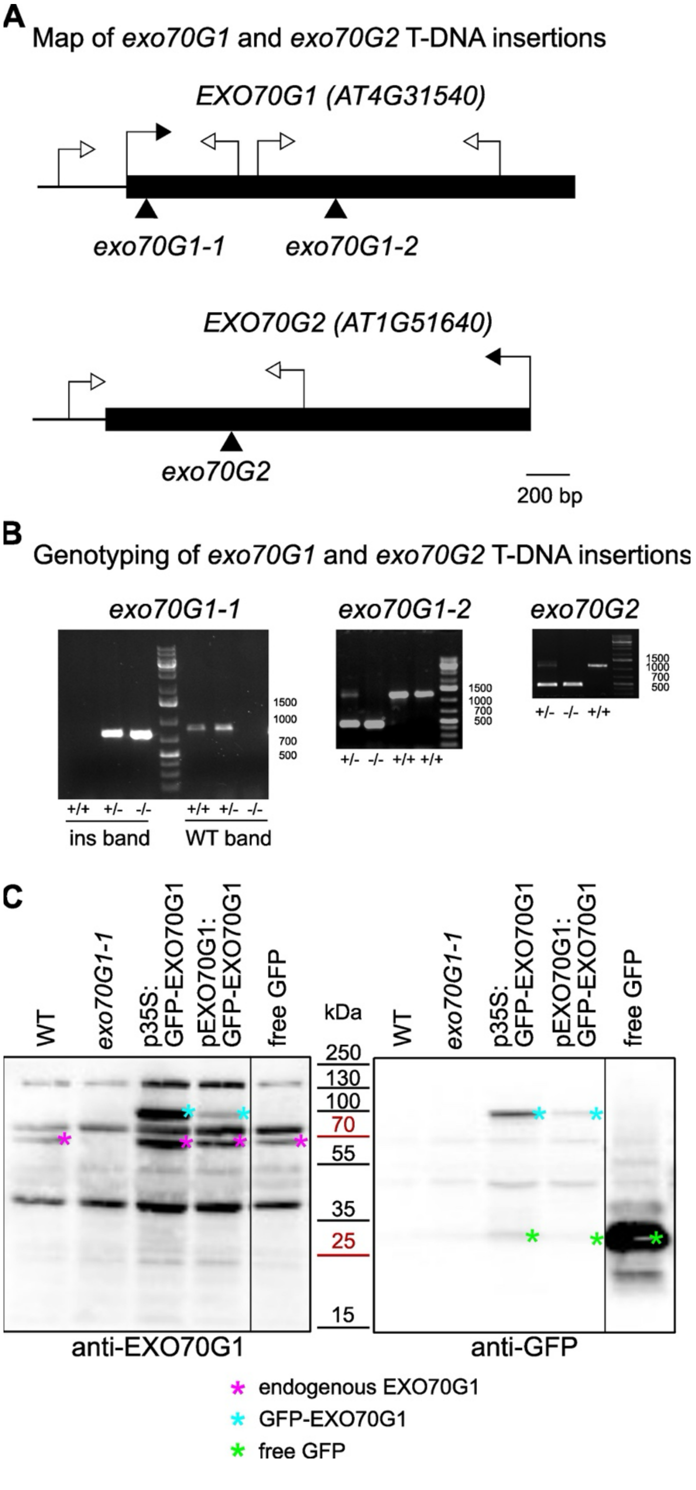


**Supplementary Figure S3.**

**Characterization of *exo70G1-1*, *exo70G1-2*, and *exo70G2* mutants. (A)** Schematic representation of T-DNA insertions in *exo70G1-1*, *exo70G1-2*, and *exo70G2*. Arrows denote positions of primers used for genotyping. **(B)** Genotyping of *exo70G1-1*, *exo70G1-2*, and *exo70G2* mutants by PCR. **(C)** Immunoblot analysis showing the presence and abundance of native and GFP-tagged EXO70G1 in the indicated mutant and transgenic lines, using polyclonal anti-EXO70G1 and anti-GFP antibodies. All transgenes were expressed in a wild-type background.

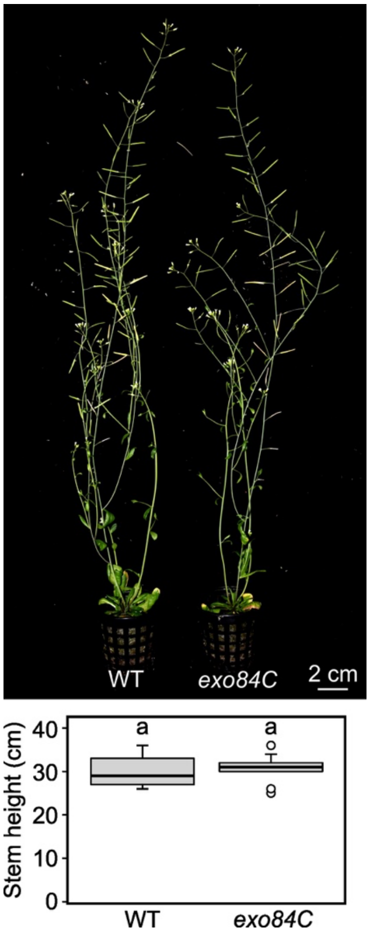


**Supplementary Figure S4.**
**Phenotype of *exo84C* mutant plants.** The representative phenotype of mature homozygous *exo84C* plants is shown alongside that of isogenic wild-type plants. No statistically significant differences in inflorescence stem height between genotypes were observed (unpaired t-test, $P > 0.05$). At least nine plants per genotype were analyzed.

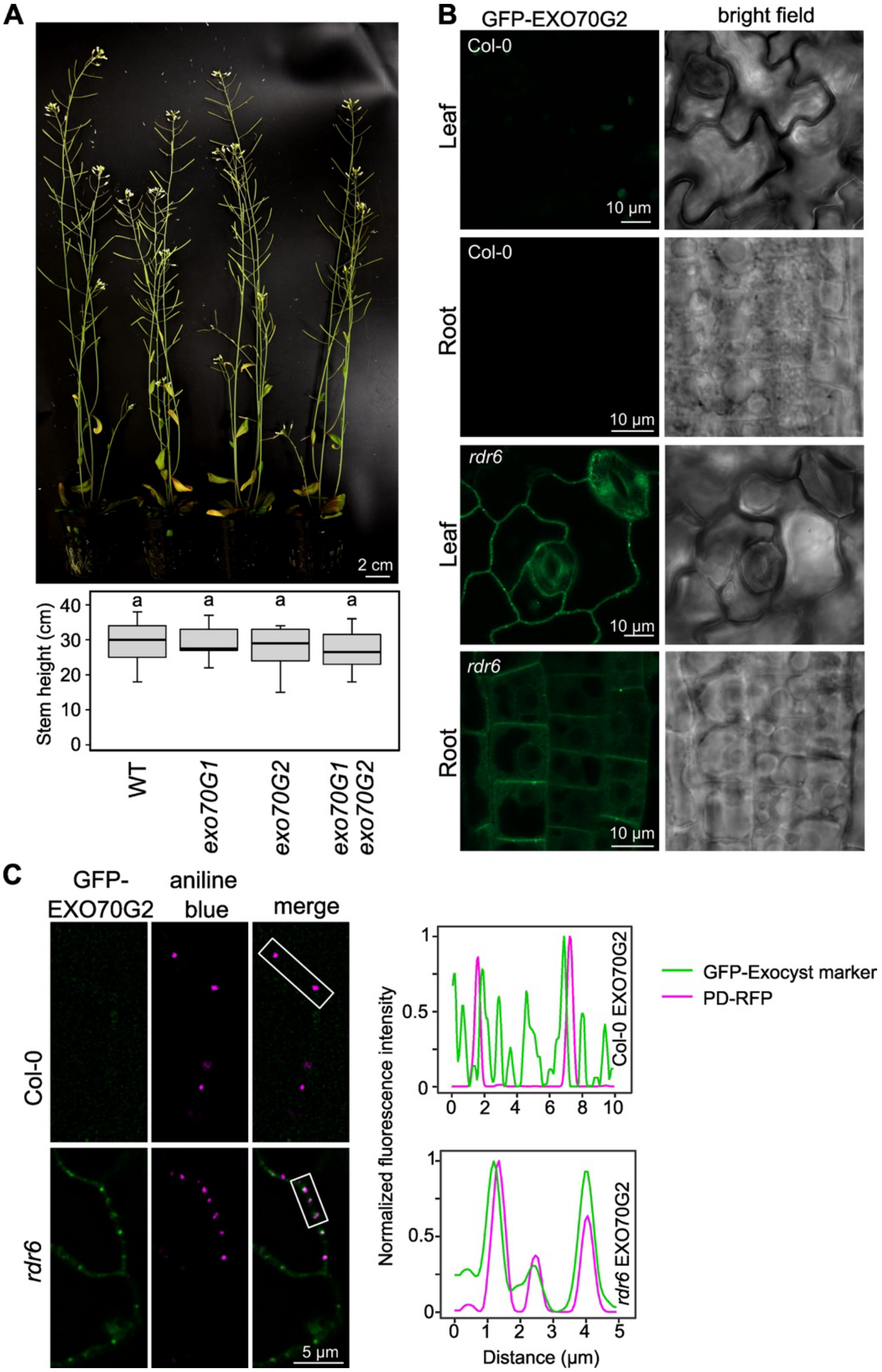


**Supplementary Figure S5.**

**Characterization of *exo70G1* and *exo70G2* mutants and EXO70G2 expression and localization. (A)** Mature homozygous single and double *exo70G* mutants shown alongside isogenic wild-type plants. Inflorescence stem height distribution is shown below; differences between genotypes were not statistically significant (Kruskal-Wallis test, $P > 0.05$). At least five plants per genotype were analyzed. **(B)** Localization of GFP-EXO70G2, expressed under the control of the 35S promoter, in Col-0 and *rdr6* genetic backgrounds in leaf and root epidermal cells. No GFP signal was detected in the Col-0 background. **(C)** Colocalization of 35S-driven GFP-EXO70G2 with aniline blue–stained callose in the leaf epidermis of Col-0 and *rdr6* plants. No GFP signal was detected in the Col-0 background.

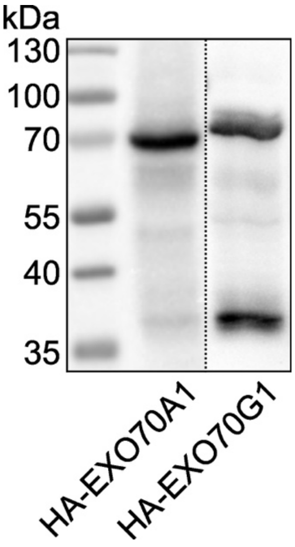


**Supplementary Figure S6.**
**Immunoblot analysis of HA-tagged EXO70A1 and EXO70G1.** *In vitro*–translated HA-tagged EXO70A1 and EXO70G1 were detected using an anti-HA antibody.

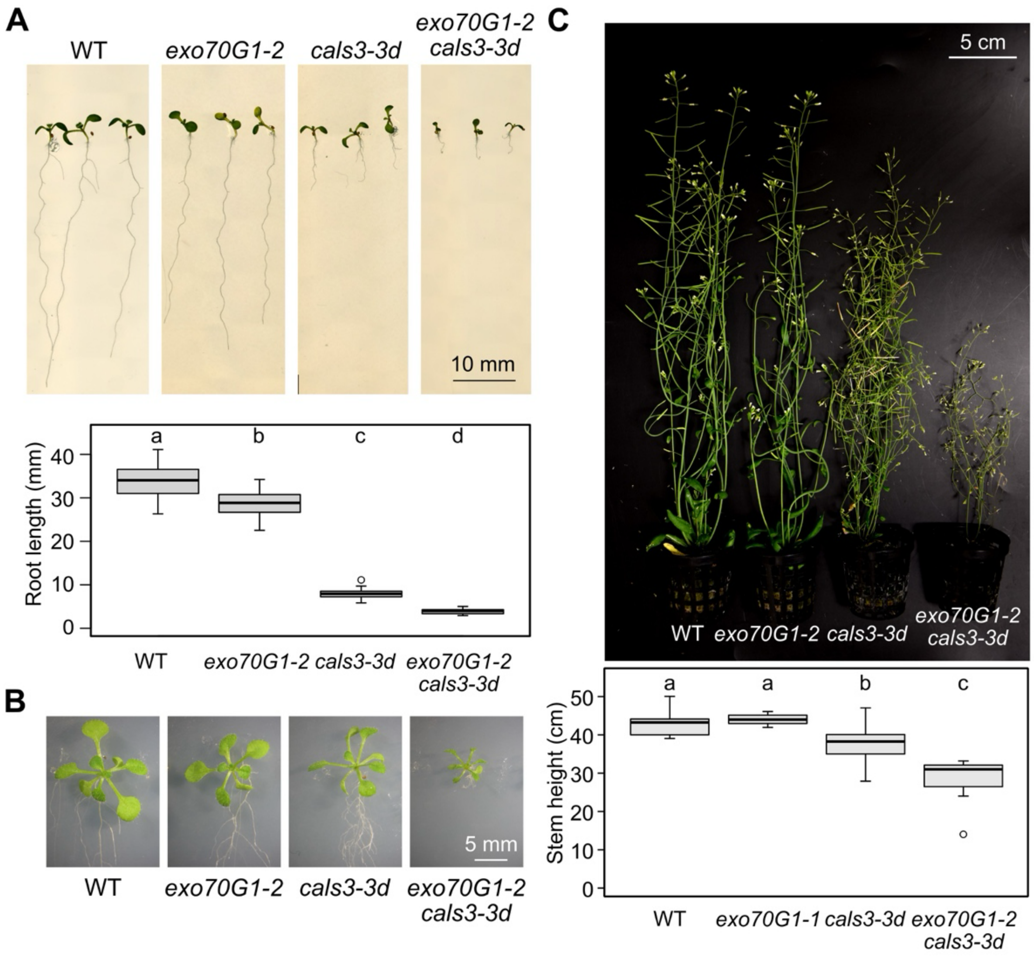


**Supplementary Figure S7.**

**Phenotype of *exo70G1-2* mutants in the *cals3-3d* background. (A)** Effects of *exo70G1-2*, *cals3-3d*, and the *cals3-3d exo70G1-2* double mutant on root length in 7-day-old seedlings. Root length was measured in at least 12 plants per genotype. Different letters indicate statistically significant differences (one-way ANOVA, $P < 0.05$). **(B)** Shoots of 14-day-old seedlings of the indicated genotypes. **(C)** Mature plants of the indicated genotypes. Inflorescence stem height was measured in at least 10 plants per genotype. Different letters indicate statistically significant differences (one-way ANOVA, $P < 0.05$).

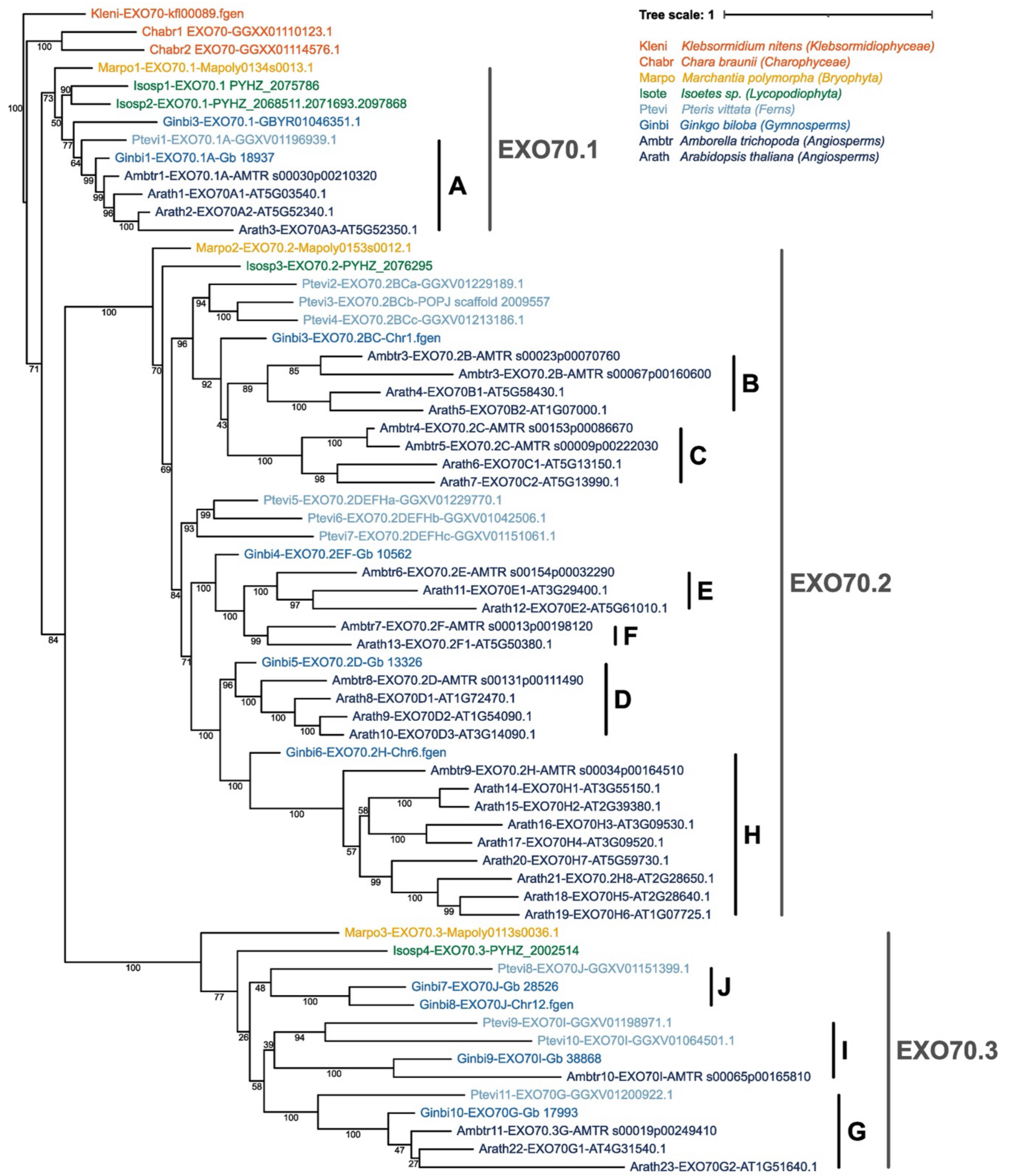


**Supplementary Figure S8.**

**Phylogenetic analysis EXO70 in streptophytes.** Protein sequences were aligned by MAFFT L-INS-I algorithm and manually trimmed. A maximum-likelihood phylogenetic tree was constructed using SeaView. Percentages of nonparametric bootstrap support calculated from 500 replicates are indicated by numbers at branches.

**Supplementary Table S1.**
**List of primers used in this study.**

| name | sequence (5' – 3') | used for | line/gene/construct |
|---|---|---|---|
| EXO70G1-1 LP | tctctggtttcggttcttgcaa | genotyping | *exo70G1-1* (SALK_099892) |
| EXO70G1-1 RP | tccacggtcattatgaaattcact | genotyping | *exo70G1-1* (SALK_099892) |
| EXO70G1-2 LP | ttgacaagatcacgagctgtg | genotyping | *exo70G1-2* (GK-209C09) |
| EXO70G1-2 RP | agttcaggaggctcttgaagg | genotyping | *exo70G1-2* (GK-209C09) |
| EXO70G2 LP | ttggtttgaccaatctcgaag | genotyping | *exo70G2* (GABI_548B11) |
| EXO70G2 RP | atttcgaggggaacaatgttc | genotyping | *exo70G2* (GABI_548B11) |
| EXO84C LP | tcaaagatcgattccgttcac | genotyping | *exo84C* (SALK_017883) |
| EXO84C RP | aagctgtttcggaaggagaag | genotyping | *exo84C* (SALK_017883) |
| LB new | gaacaacactcaaccctatctcgggc | genotyping | SALK lines |
| GK8409 | atattgaccatcatactcattgc | genotyping | GABI-Kat lines |
| AtEXO84C-F_EcoRI | atagaattcatggagagcagcgaggaag | cloning | AD-, BD-AtEXO84C-N |
| AtEXO84C-N-S-R_SalI | atagtcgactcactcatcagtttcattcaagtcaaag | cloning | AD-, BD-AtEXO84C-N |
| AtEXO84C-C-F_XmaI | atacccgggaggcttggaatcaccctctg | cloning | AD-EXO84C-C |
| AtEXO84C-F_XmaI | atacccgggaatggagagcagcgaggaag | cloning | AD-, BD-EXO84C |
| AtEXO84C-S-R_SalI | atagtcgactcaagactcagaatcggtgaaag | cloning | AD-, BD-EXO84C, AD-EXO84C-C |
| pAtEXO84C-F_KpnI | ataggtaccatcgtcatcatgcgtttcttttg | cloning | pAtEXO84C::AtEXO84C in pENTR3C |
| AtEXO84C-R-NS_NotI | atagcggccgcaaagactcagaatcggtgaaagatg | cloning | pAtEXO84C::AtEXO84C in pENTR3C |
| pAtEXO70G1-F_attB4 | ggggacaactttgtatagaaaagttgctgtcaaagttgtcatttgt | cloning | pAtEXO70G1 in pDONR P4-P1r |
| pAtEXO70G1-R_attB1r | ggggactgctttttgtacaaacttgctaaattcaatcaaaataaaac | cloning | pAtEXO70G1 in pDONR P4-P1r |
| AtEXO70G1-F_attB2r | ggggacagctttcttgtacaaagtggctatgatggcggagtctac | cloning | AtEXO70G1 in pDONR P2r-P3 |
| AtEXO70G1-R_attB3 | ggggacaactttgtataataaagttgctcagacaactgcagaagtag | cloning | AtEXO70G1 in pDONR P2r-P3 |
| AtEXO70G2-F_attB2r | ggggacagctttcttgtacaaagtggctatggctgaatcaaagagact | cloning | AtEXO70G2 in pDONR P2r-P3 |
| AtEXO70G2-R_attB3 | ggggacaactttgtataataaagttgctcatatagcttctaatgtcgag | cloning | AtEXO70G2 in pDONR P2r-P3 |
| AtEXO70A1-F | gatgttccagattacgctatggctgttgatagcaga | cloning | HA-AtEXO70A1 |
| AtEXO70A1-S-R_NotI | atagcggccgcttaccggcgtggttc | cloning | HA-AtEXO70A1 |
| AtEXO70G1-F | gatgttccagattacgctatgatggcggagtctact | cloning | HA-AtEXO70G1 |
| AtEXO70G1-S-R_NotI | tatgcggccgctgcaggtcgactcagacaactgcagaagtag | cloning | HA-AtEXO70G1 |
| pTNT-HA-oh_SalI | aagtcgacgccgccaccatgtacccatacgatgttccagattacgctatg | cloning | HA-AtEXO70A1 |
| pTNT-HA-oh_XhoI | atactcgaggccgccaccatgtacccatacgatgttccagattacgctatg | cloning | HA-AtEXO70G1 |
